# Multilayer-assembled microfluidic chip with Tesla-valve channels: Combined biochemical gradient and mechanical stiffness interface for 3D SKOV3 cell culture

**DOI:** 10.64898/2026.09.15.751867

**Authors:** Xiaolu Zhu, Wei Qian, Hao Cheng

**Author notes:** Correspondence: Xiaolu Zhu (X. Zhu). These authors contributed equally to this work (Xiaolu Zhu, Wei Qian).

## Abstract

Biochemical gradients and mechanical microenvironments synergistically regulate tumor cell proliferation, migration, and phenotypic evolution, which are critical for recapitulating *in vitro* tumor physiological microenvironments. In this work, a multilayer-assembled microfluidic chip integrated with Tesla-valve flow channels was developed for 3D SKOV3 cell culture under combined biochemical gradients and mechanical stiffness interfaces. The reconfigurable multilayer PMMA structure enables flexible construction of diverse concentration gradient fields, while the embedded Tesla-valve geometry effectively stabilizes fluid flow, reduces flow velocity fluctuation, and generates smoother, more stable biochemical gradients compared with conventional curved channel designs. Multiphysics simulations were performed to systematically verify the flow field distribution and gradient formation performance of the optimized channel structure. Furthermore, by constructing heterogeneous hydrogel microenvironments consisting of rigid Pluronic F127 and soft dextran hydrogel inside microfluidic chambers, stable mechanical stiffness interfaces were successfully fabricated. Combined with TGF-β1 biochemical stimulation, the platform was applied to explore the durotactic migration and morphological changes of 3D-cultured SKOV3 cells at stiffness interfaces. Fluorescence staining results demonstrated that mechanical boundary conditions significantly affected cell migration behavior and cell aggregation phenotypes under consistent biochemical induction. With the advantages of low cost, simple assembly, good biocompatibility, and controllable dual physicochemical microenvironments, this multilayer microfluidic platform provides a reliable and efficient strategy for *in vitro* tumor microenvironment simulation and tumor cell mechanobiology research.

## 1. Introduction

Microfluidic chip technology has been widely adopted for anticancer drug-screening applications, including nanoparticle-based, paper-based and polymer-based platforms^[1, 2]^. Using soft lithography, microchannels, microgrooves, and culture chambers can be patterned on PDMS to fabricate high-performance microfluidic devices rapidly. Such platforms support both two-dimensional (2D) and three-dimensional (3D) cell culture^[1]^. They can also supply continuous culture medium and provide micro-reaction compartments for testing cellular responses to drugs, toxicants, or other chemical stimuli^[3, 4]^.

Nanoparticle-integrated microfluidic architectures have been exploited for chemotherapeutic drug screening, where embedded microstructures enhance fluid mixing and drug-cell interactions; one such system evaluated methotrexate against osteosarcoma cells and demonstrated the therapeutic potential of nanoparticle-based cancer interventions^[2]^. Paper-based microfluidics represent another promising branch of this technology. A two-layer paper device separates cell culture compartments from underlying self-driven perfusion channels, enabling direct cell culture on hydrophilic substrates without hydrogel embedding [3]. By combining microfluidic manipulation with microelectrodes within a 3D tumor-mimicking niche, the platform permits accurate, real-time readouts of cancer cell responses to pharmacological cues and supports anticancer drug-screening workflows^[3]^.

Other polymer-based microfluidic chips, such as the PMMA-SU8-glass based microfluidic platform with electrical-response monitoring also represents an alternative strategy for on-chip drug assays. One microfluidic system integrated with microsensors records electrical signals from cancer cells embedded inside 3D hydrogels upon chemotherapeutic challenge^[4]^. It has been applied to assess dose-dependent effects of carboplatin on B16-F10 and 4T1 cell lines, as well as paclitaxel responses in prostate cancer cells. Nevertheless, the device possesses a relatively simple layout. Drug-laden streams travel counter-directionally along an offset parallel path adjacent to cell-laden hydrogel within the reaction zone. Even under this side-by-side flow configuration, interfacial shear stress is still generated, and only a single drug concentration can be interrogated per experimental run^[4]^. Further, Shirure et al.^[5]^ reported a PDMS-based microfluidic platform that recapitulates vascular-mediated drug delivery to tumor cell populations. This device mimics biological mass transport occurring at arterial capillary ends within the tumor microenvironment. Nevertheless, the generated gradient is confined to a relatively narrow spatial region, limiting its capacity to examine cellular responses across broad gradient ranges.

The therapeutic potency of single-agent chemotherapy remains limited, and combinatorial drug regimens often yield superior outcomes for cancer treatment. Shen et al.^[6]^ assessed monotherapy and combination effects of two chemotherapeutic agents, doxorubicin and cisplatin, against MCF-7 and HepG2 tumor cell lines. They fabricated a PDMS-based microfluidic device equipped with specially designed triple microchannel architectures to establish concentration gradients spanning multiple discrete cell-culture chambers. Benefiting from the double-spiral micromixer which also acts as a fluid damper, the generated concentration profiles remain stable across a broad flow-rate window ranging from 0.1 to 200 μL min⁻¹, rendering this platform well-suited for evaluating tumor responses under multi-drug combinatorial interventions. Zhang et al. ^[7]^ reported a high-throughput microfluidic screening platform equipped with Christmas-tree-style mixers to produce logarithmic concentration gradients for pairwise drug combinations. Benefiting from its three-layer PDMS architecture, the platform enabled multiplexed combinatorial drug assessment on tumor spheroids, featuring low sample consumption and rapid readout for precision-oncology screening applications. Nevertheless, this monolithic PDMS device relies on irreversible plasma bonding, where channel layouts are fixed after fabrication and cannot be reconfigured to generate alternative gradient profiles. In addition, drug delivery is implemented by continuous perfusion, which exposes 3D tumor spheroids to persistent fluid-induced shear stress; the bonded structure also prevents disassembly for thorough cleaning, restricting the chip to single-use only. Khoo et al. ^[8]^ designed a multi-channel microfluidic device with adjustable valves for exploring the combined effect of doxorubicin and aspirin on circulating tumor cells, which supports 10 different drug concentrations but only provides fixed single-concentration drug stimulation for samples, failing to form continuous chemical concentration gradients. Moreover, existing automated valve-equipped microfluidic devices are mostly tailored for fixed cell culture positions and single functional modes, lacking structural reconfigurability.

Recent advanced microfluidic systems have further promoted the development of gradient-based cell culture and drug screening. Some integrated microfluidic platforms have realized stable linear chemical gradient generation and long-term dynamic cell culture, effectively simulating the tumor biochemical microenvironment for preclinical drug evaluation ^[9]^. Nevertheless, most integrated gradient chips rely on permanently bonded microchannel structures with fixed gradient modes, which cannot be flexibly adjusted to produce multiple gradient profiles for comparative biological research. In addition, scalable microfluidic screening platforms based on 3D microtumor models have been proven to support high-throughput multi-drug combination screening, significantly improving the efficiency of precision drug screening^[10]^. However, such high-throughput devices usually require sophisticated microfabrication and external auxiliary control systems, resulting in high cost and poor accessibility for conventional laboratory applications. Most recently, improved multi-functional microfluidic platforms have been proposed for simultaneous multi-drug screening and 3D cell culture ^[11]^, but these devices still adopt traditional perfusion-driven mass transfer modes, which inevitably introduce undesirable fluid shear stress on 3D cultured cells and impair cell physiological activity, limiting their application in high-precision gradient cell research.

Another notable study reported a dual-channel, three-layer PDMS microfluidic chip for combinatorial drug screening^[12]^. Equipped with serpentine microchannels, this device generates a stable one-dimensional stepwise linear concentration gradient spanning six culture chambers. CAL-27 cell experiments reproduced dose-dependent cytotoxic responses consistent with conventional well-plate results, enabling full-range gradient measurement on a single chip without preparing multiple discrete samples. Even so, this irreversibly bonded, monolithic PDMS chip lacks structural reconfigurability and only supports a fixed gradient profile. Continuous perfusion delivery inevitably introduces fluid shear stress on cultured cells, and its non-detachable structure restricts the device to single-use applications. Similarly, a high-throughput three-layer PDMS microfluidic chip has been developed to generate logarithmic stepwise drug concentration gradients for combinatorial anticancer drug screening^[13]^. Although it enables parallel assays of 3D tumor spheroids and patient-derived tumor cells for precision oncology evaluation, this device still suffers from fixed channel geometry, perfusion-induced shear stress, and single-use limitations.

Despite the significant progress of current microfluidic technology in 3D cell culture and gradient-based tumor biological research, existing microfluidic devices still suffer from inherent structural and functional limitations that restrict their practical application. First, most conventional microfluidic chips are fabricated via customized soft lithography and irreversible bonding processes. The integrated fixed structure leads to poor reconfigurability: each chip can only generate a single fixed concentration gradient mode, and structural modification or multi-gradient generation requires repeated, complex fabrication procedures involving mold preparation, glue pouring, curing, and bonding, which is time-consuming, costly, and inflexible for diverse experimental requirements. Second, traditional gradient generation structures generally adopt direct perfusion of drug solution into cell culture chambers, and the fluid convection inevitably produces shear force that directly acts on cultured cells, resulting in cell damage and abnormal growth states, which severely interferes with the accuracy of cell response detection under chemical gradients. Third, mainstream PDMS-based chips are difficult to disassemble and clean after experiments, causing residual cell and reagent contamination and poor reusability, further limiting their application scalability in long-term biological research. Beyond biochemical gradients, tumor progression is also modulated by mechanical cues such as matrix stiffness^[14–17]^. Most existing gradient microfluidic platforms only establish biochemical gradients, and few systems can simultaneously construct tunable biochemical gradients and defined mechanical stiffness interfaces to mimic the heterogeneous tumor microenvironment, which limits the investigation of combined biochemical and mechanical regulation of tumor cell behavior.

To address the above critical technical bottlenecks, this work develops a novel disassembled, reconfigurable multi-layer assembled microfluidic chip for high-stability horizontal concentration gradient generation and 3D cell culture. Distinct from traditional monolithic fixed-structure microfluidic devices, the proposed chip adopts a modular layered assembly architecture consisting of detachable cover plates, multi-functional flow channel layers, and a limiting base. The template-based layered structure enables free replacement of flow channel layers with different configurations, which can rapidly generate four distinct stable horizontal chemical concentration gradients without repeated chip fabrication, greatly simplifying the manufacturing process and reducing experimental costs. Benefiting from the specially designed narrow microchannel structure between the flow channel layer and the culture chamber, bioactive factors diffuse into the 3D cell culture chamber through passive diffusion rather than direct fluid perfusion, which effectively eliminates fluid shear force interference on cell growth and maintains the physiological activity of cultured cells. In addition, the fully detachable assembly design facilitates thorough cleaning and repeated reuse of the chip, while the low-cost PMMA and silicone membrane materials further improve the accessibility and practicability of the device. A Tesla-valve flow channel design was proposed and integrated to stabilize the flow field and achieve smoother and more uniform concentration gradient distribution. Furthermore, heterogeneous hydrogel constructs were embedded within the chambers to form mechanical stiffness interfaces, enabling the investigation of tumor cell behaviors under combined biochemical and mechanical microenvironmental stimuli. The biocompatibility and application performance of the chip are validated via SKOV3 cell culture experiments, providing a cost-effective, flexible, and high-precision technical platform for horizontal concentration gradient-mediated tumor cell research and anticancer drug screening.

## 2. Model description and working principle

Figure 1 presents the structural schematic of the multilayer assembled microfluidic chip, which comprises five sequentially stacked functional layers. As illustrated in the exploded and assembled views (Fig. 1a–1d), the entire device measures 40 mm × 40 mm and consists of a top PMMA cover plate (Layer 1), a first silicone sealing membrane (Layer 2), a patterned PMMA channel substrate (Layer 3), a second silicone sealing membrane (Layer 4), and a bottom transparent glass substrate (Layer 5).

**Fig. 1.**
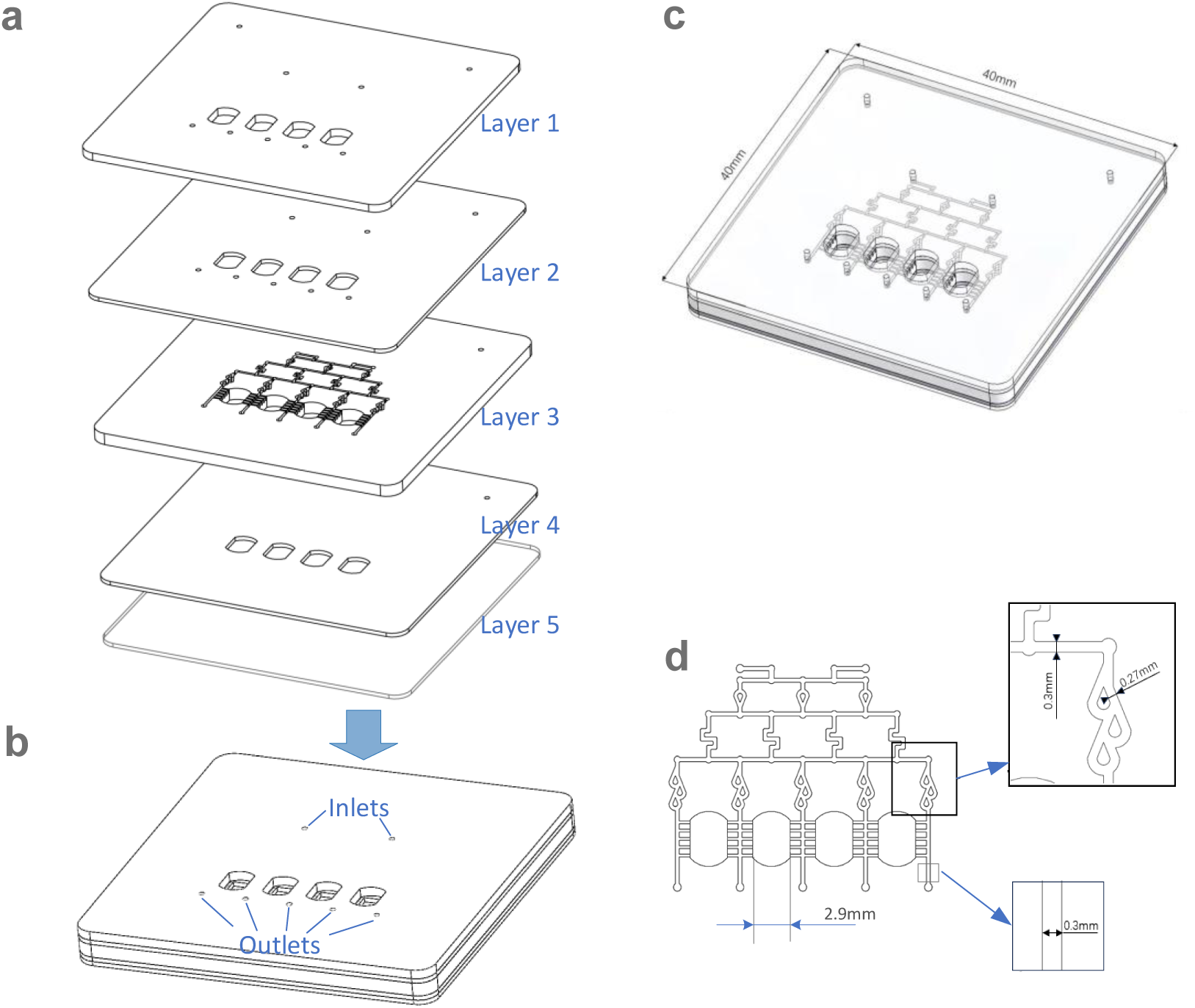
Schematic structural design of the multi-layer assembled microfluidic chip. (a) Exploded-view schematic; (b) assembled configuration with an arrow illustrating the stacking-assembly process; (c) transparent wire-frame view of the assembled device annotated with overall dimensions (40 mm × 40 mm), revealing the internal layout of microchannels; (d) dimensional drawing of local structures. The chip is stacked sequentially by five components: a top PMMA cover (Layer 1), a first silicone sealing membrane (Layer 2), a patterned PMMA flow-channel substrate (Layer 3), a second silicone sealing membrane (Layer 4), and a bottom glass substrate (Layer 5).

The top PMMA cover plate is fabricated with two locating holes, two medium inlets, five outlets, and four independent culture chambers. The overlying silicone membrane replicates the same structural layout to ensure structural alignment and tight sealing. The middle PMMA layer serves as the core functional channel layer, retaining identical chamber and port configurations while integrating gradient-generating microchannels to recapitulate physiological biochemical microenvironments. Culture medium flows through these microchannels and steadily enters the downstream culture chambers. The bottom silicone membrane further ensures interface sealing and structural consistency, whereas the underlying optical glass substrate provides high transparency for real-time microscopic observation of cellular behavior.

The consistent inlet and outlet positioning across the upper three layers enables smooth and stable fluid delivery throughout the microchannel network. The double-layer silicone membranes effectively eliminate medium leakage and ensure reliable chip assembly. Narrow microchannels embedded in the PMMA substrate reduce flow velocity, thereby minimizing fluid shear stress on encapsulated cells and allowing exclusive investigation of biochemical gradient effects. Combined with precisely controlled syringe pump injection, this design generates four distinct stable drug concentrations within the four chambers.

Notably, the replaceable patterned channel layer enables flexible adjustment of gradient sequences without reconstructing the entire device, substantially reducing fabrication cost and experimental cycle time. Benefiting from the optically transparent architecture, the chip supports real-time microscopic imaging and quantitative image analysis, permitting rapid and accurate evaluation of drug-induced cytotoxicity, morphological changes, and migration behaviors of cancer cells. Compared with conventional PDMS–glass microfluidic platforms, the proposed multilayer assembly strategy consumes fewer materials and reagents. It also avoids repeated fabrication, alignment, and cleaning of multiple customized chips, thereby simplifying experimental operations and reducing potential operational errors.

## 3. Numerical simulation

Different types of concentration gradients can be realized on the presented microfluidic chip by replacing the third-layer channel substrate. Two distinct 3D channel geometries of this reconfigurable chip were built using SolidWorks (Fig. 2a, Fig. 3a-b). Given the well-established applicability of the finite-element method (FEM) for microfluidic simulations, the geometries were imported into COMSOL Multiphysics 5.5 (trial version) to conduct numerical analysis and guide chip design.

**Fig. 2.**
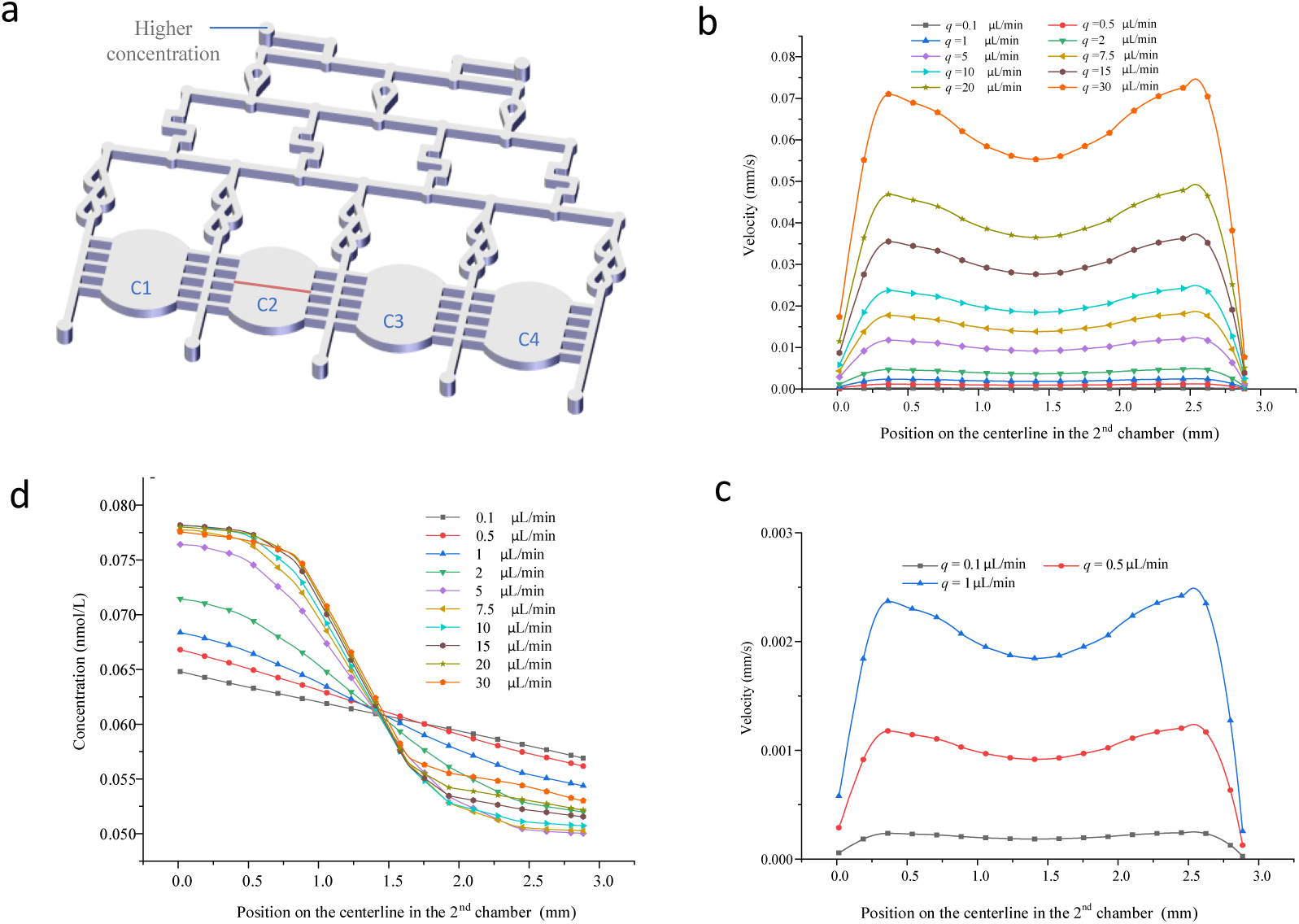
Multiphysics simulation of flow velocity and solute concentration within the culture chambers of Layer 3. (a) Geometric model of the computational domain. (b) Velocity profiles along the central sampling line (red line in panel a) of the 2nd culture chamber under varying inlet flow rates. (c) Magnified velocity curves for low-flow-rate conditions (*q* = 0.1, 0.5 and 1 μL/min). (d) Solute-concentration distribution along the centerline of the 2nd culture chamber.

**Fig. 3.**
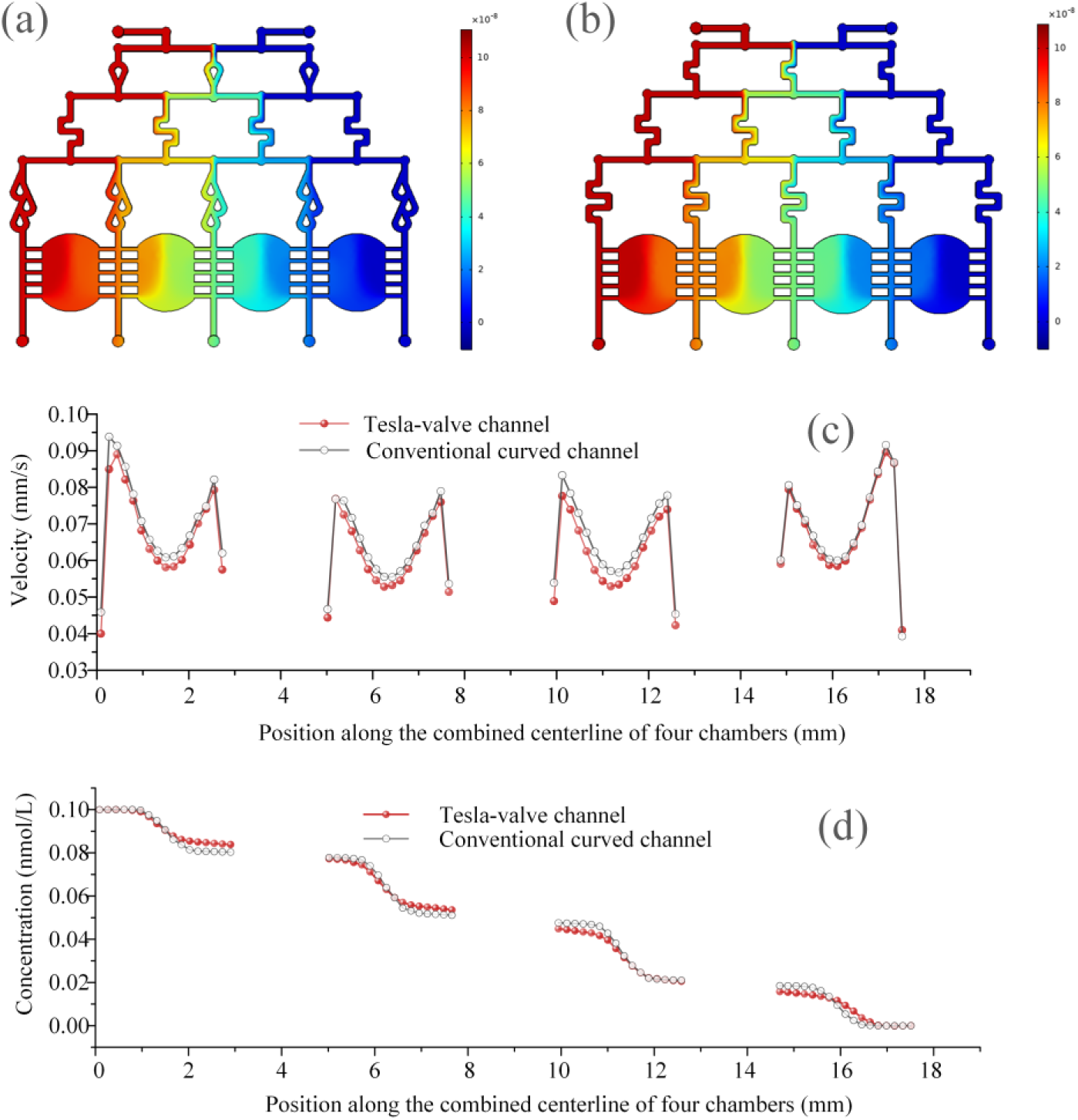
Numerical simulation comparison between conventional curved-channel and Tesla-valve-integrated microfluidic configurations. (a, b) Simulated concentration contour plots of (a) Tesla-valve channel within the multi-layer microfluidic device and (b) conventional curved channel. (c) Velocity profiles extracted along the combined centerline across four sequential culture chambers. Local velocity peaks correspond to geometric junctions connecting culture chambers and horizontal narrow connecting channels. (d) Concentration profiles obtained along the combined centerline for the two channel designs. Simulations in this figure were performed at an inlet flow rate of 30 μL/min to exaggerate fluid-field differences for mechanistic interpretation.

### 3.1 Multiphysics coupled mechanism

The fluid domain of the microfluidic model consists of two inlets, gradient-generating channels, narrow connecting channels, four cell culture chambers, and five outlets. Material assignments for chip components were selected from COMSOL’s built-in material library, with material parameters configured to match the physical properties of real-world chip constituents. Default library parameters were adopted for most solid components, whereas custom parameters were defined for the hydrogel domain: density = 1000 kg/m³, dynamic viscosity = 2 × 10⁻³ Pa·s, and porosity = 0.8.

Two physical interfaces were activated for simulation: Creeping Flow and Transport of Diluted Species in Porous Media, accounting for fluid hydrodynamics and solute mass-transport behavior, respectively.

#### 3.1.1 Creeping Flow

Low-Reynolds-number conditions dominate inside microchannels, and the fluid exhibits laminar flow characteristics. Flow fields throughout the device were solved using the Creeping Flow interface, which is governed by the continuity equation and Navier-Stokes equations:

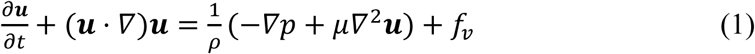

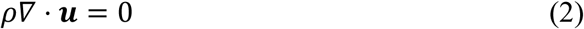

where *ρ* denotes fluid density, **u** is the velocity vector, *µ* represents dynamic viscosity, ∇*p* is the pressure gradient, and *f_v_* refers to body force per unit fluid mass. Solving Equations (1) and (2) over the computational domain yields the full velocity-field distribution within microchannels as well as global pressure profiles across the system. For creeping–flow regimes, inertial terms in Eq. (1) can be safely neglected.

#### 3.1.2 Transport of diluted species in porous media

This microfluidic chip is designed for 3D cell culture and investigation of cell behaviors under varied biochemical concentration gradients. The *Transport of Diluted Species in Porous Media* interface was adopted to simulate solute-drug transport within both free-flow channels and porous hydrogel domains. Mass transport within the model is governed by the convection-diffusion equation:

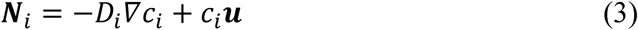

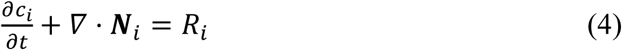

where ***N****_i_* is the molar flux (mol/(m^2^·s)), *D_i_* denotes the diffusion coefficient (m^2^/s), *c_i_* represents solute concentration (mol/m^3^), ***u*** is the velocity vector (m/s), and *R_i_* stands for the species reaction rate (mol/(m^3^·s)), where subscript *i* denotes the *i*-th solute species. In the present simulation, solute consumption or chemical reactions were neglected; therefore, *R_i_* = 0.

### 3.2 Boundary conditions

The numerical model couples the *Creeping Flow* and *Transport of Diluted Species in Porous Media* interfaces. For the *Creeping Flow* physics, the upper cylindrical ports were defined as flow-field inlets, whereas the lower ports served as outlets. Within the solute-transport interface, one upper-left cylindrical inlet was assigned as the drug-loading inlet, and the opposite upper-right inlet supplied drug-free medium. Specified-flow-rate boundary conditions were imposed on the inlets, and zero-pressure conditions were applied at all outlets. The four culture-chamber domains were assigned hydrogel material properties, and microchannel regions were defined as aqueous medium. The whole computational domain, covering microchannels and culture chambers, was discretized using COMSOL’s built-in physics-controlled meshing algorithm. The mesh resolution was set to *finer* to achieve mesh-independent numerical results. The final mesh contained 1931302 domain elements, 198189 boundary elements, and 16506 edge elements.

## 4. Materials and methods

### 4.1 Fabrication of each layer of the microfluidic chip

The multilayer microfluidic chip consists of five stacked functional layers: a top PMMA cover (Layer 1), a first silicone sealing film (Layer 2), a PMMA flow-channel substrate (Layer 3), a second silicone sealing film (Layer 4), and a bottom optical glass substrate (Layer 5). All layers have a uniform planar dimension of 40 mm × 40 mm, with thicknesses of 1.5 mm, 0.5 mm, 1.5 mm, 0.5 mm, and 0.5 mm for Layer 1 to Layer 5, respectively, as shown in Fig. 1.

Layer 1 was fabricated from polymethyl methacrylate (PMMA) via CO₂ laser cutting. Layers 2 and 4 were patterned on silicone films using ultraviolet picosecond laser cutting. The core PMMA flow-channel layer (Layer 3) was processed by computer numerical control (CNC) machining and further treated with hydrogen peroxide under hydrothermal conditions for surface optimization. The bottom optical glass substrate (Layer 5, DEEN OPTICS) offers high transparency to facilitate microscopic observation of cells.

Layer 1 and Layer 2 share identical through-hole layouts, including two positioning holes, two fluid inlets, four cell culture chambers, and five fluid outlets. All inlet, outlet, and positioning holes have a uniform radius of 0.6 mm. Each culture chamber is structurally composed of 2 mm-radius arc ends and 2.75 mm straight side edges. As the gradient-generating core structure, Layer 3 contains two positioning holes and integrated microchannels with a consistent width of 0.3 mm and depth of 0.5 mm. Differing from Layer 2, Layer 4 only reserves positioning holes and chamber cutouts without inlet and outlet channels. Stainless-steel capillary tubes were fixed at all channel inlets and outlets using epoxy adhesive. Before assembly, all chip components were thoroughly cleaned with 75% ethanol. Ultrasonic cleaning was performed for 5 min at 30 °C. The cleaned components were naturally dried inside a Class II biosafety cabinet to ensure sterile and dust-free conditions, followed by vacuum drying at 80 °C for 1 h for further dehydration and sealing preparation.

### 4.2 Cell culture and pigment test

Human SKOV3 cells were purchased from Cobioer Co., Ltd., Nanjing, China. Cells were cultured in RPMI-1640 medium supplemented with 10% (v/v) fetal bovine serum (Cat. No.: 13011-8611, Every Green, Zhejiang Tianhang Biotechnology Co., Ltd., China) and 1% (v/v) penicillin–streptomycin (Cat. No.: SV30010, HyClone, Logan, UT, USA). Cells were incubated at 37 °C with 5% CO₂. The culture medium was refreshed every two days, and cells were routinely passaged when reaching approximately 85% confluence.

Red and blue dyes were used to visually verify the concentration gradient generation capability of the microfluidic chip.

### 4.3 Preparation of the microfluidic chip for cell-seeded hydrogels

To ensure biosafety and experimental reproducibility, all chip components were cleaned with ethanol and rinsed with PBS to remove residual alcohol, followed by air-drying in a Class II biosafety cabinet. After assembly, the microfluidic chip was placed in a vacuum drying oven to eliminate residual air and enhance interfacial sealing. Prior to cell experiments, 1 mL syringes filled with complete culture medium were connected to stainless-steel capillary inlets through transparent silicone tubing. The fully assembled chip was further sterilized via ultraviolet irradiation for 30 min.

SKOV3 cells were encapsulated in dextran-based hydrogels at a final density of 2000 cells/µL. The molar concentrations of maleimide groups, RGD peptides, and CD-linkers were set to 3 mM, 11.25 μM, and 2 mM, respectively. Detailed proportions of the hydrogel precursor components are summarized in Table 1. For each culture chamber, CD-linker solution was pre-added first. The cell-laden hydrogel precursor solution (9 μL per chamber) was aspirated using a RAININ Pos-D pipette and gently mixed with the preloaded CD-linker solution to initiate gelation, with strict precautions taken to prevent bubble formation.

**Table 1.** Composition of cell-laden dextran hydrogel.

| Component name | Volume / $\mu\text{L}$ |
| --- | --- |
| Maleimide-Dextran ($C_{\text{Mal}}$: 30 mmol/L) | 4 |
| CD-Linker ($C_{\text{SH}}$: 20 mmol/L) | $1 \times 4$ |
| Thioglycerol (–SH group: 0.3 mmol/L) | 12 |
| RGD peptide (–SH group: 0.3 mmol/L) | 1.5 |
| 10-fold CB (pH = 5.5) | 2.5 |
| Water | 10 |
| RPMI-1640 (with cells) | 6 |

*Note: $1 \times 4$ indicates 1 $\mu\text{L}$ of CD-Linker solution added to each of the four culture chambers.*

Pluronic F127-based hydrogel (SunP Gel S1, SunP Biotech Co., Ltd., Beijing, China) was used to surround the inner dextran hydrogel for fabricating mechanical stiffness interfaces within the microfluidic chambers. The Young’s modulus of Pluronic F127 hydrogel ranges from several thousand to 10 000 Pa, whereas the dextran hydrogel adopted in this study had a modulus below 1 kPa. Qualitative mechanical assessment by spatula contact showed that the Pluronic F127 hydrogel exhibited much less deformation, confirming its substantially higher stiffness compared with the dextran hydrogel.

After gelation was complete, fresh culture medium was overlaid on the hydrogels. Culture medium was continuously perfused into each culture chamber at 0.3–0.5 μL/min to minimize fluid shear stress on the encapsulated cells. Under continuous perfusion, TGF-β1 in the medium formed stable biochemical concentration gradients across the hydrogel and diffused toward the embedded SKOV3 cells. Periodic time-lapse imaging was carried out to record cellular phenotypes within the microfluidic chambers.

### 4.4 Live/Dead assay

Cell viability was evaluated using the LIVE/DEAD Viability/Cytotoxicity Kit (Cat. No.: L3224, Thermo Fisher Scientific, Eugene, OR, USA) after 24 h of cell culture within the microfluidic chip. Samples were rinsed twice with phosphate-buffered saline (PBS), with 3 min for each wash. Subsequently, 40 µL of Live/Dead working solution (3.6 µM ethidium homodimer-1 and 2.8 µM calcein-AM dissolved in DME/F-12 medium) was introduced into each culture chamber. Samples were incubated at 37 °C under a 5% CO₂ atmosphere for 25 min prior to imaging. Fluorescence signals were captured using an inverted fluorescence microscope.

### 4.5 F-actin and nucleus staining

For cytoskeletal and nuclear staining, cells were rinsed three times with PBS (5 min per wash) and fixed with 4% formaldehyde (Cat. No.: 28908, ThermoFisher, Rockford, IL, USA) at room temperature for 30 min. After fixation, samples were rinsed with PBS three more times (5 min each). Cell permeabilization was carried out using 0.1% Triton X-100 (Cat. No.: X100-5ML, SIGMA-ALDRICH, St. Louis, MO, USA) in PBS for 2 min, followed by three additional PBS washes.

Alexa Fluor 488 phalloidin (Cat. No.: A12379, Invitrogen, ThermoFisher, Rockford, IL, USA) powder was dissolved in DMSO (Cat. No.: 276855, SIGMA-ALDRICH, St. Louis, MO, USA) to prepare the stock solution following the manufacturer’s instructions. The working solution was freshly prepared by diluting 1 µL of stock solution into 1000 µL PBS. Samples were incubated with the phalloidin working solution for 45 min at room temperature in the dark to label F-actin filaments.

For nuclear staining, DAPI stock solution (Cat. No.: 62247, ThermoScientific, Dreieich, Germany) was diluted with PBS to a final concentration of 138 ng/mL. Each chamber was loaded with 30 µL of diluted DAPI solution and incubated for 10 min at room temperature in the dark. Finally, all samples were washed three times with PBS (5 min per wash) to remove residual stain prior to imaging.

## 5. Results

### 5.1 Simulation results for the flow velocity and substance concentration inside the chamber on the chip

Flow-field and solute-transport behaviors within PMMA microchannels were simulated in COMSOL Multiphysics by coupling the *Creeping Flow* and *Transport of Diluted Species* interfaces. Inlet flow rates varied from 0.1 μL/min to 30 μL/min, and the inlet solute concentration was fixed at 1×10⁻⁷ mol/m³. Cross-sectional sampling lines were positioned across the central region of the second culture chamber to quantify spatial distributions of flow velocity and solute concentration.

Under constant-inlet-flow-rate conditions, the internal flow velocity inside the chamber decreased as PMMA channel depth increased (Fig. S2). At an inlet flow rate of 30 µL/min, the chamber-interior flow velocity obtained for a channel depth of 0.5 mm was markedly lower than those for 0.3 mm and 0.4 mm channel depths. Higher flow velocity, and consequently elevated shear stress, occurred near both ends of the chamber relative to its central area.

Channel-and-chamber depth (*d*) directly modulates flow-velocity magnitude. In this work, channel depth was evaluated within 0.1–0.5 mm, taking multiple practical constraints into account: minimal consumption of reagents and hydrogel materials, physical space requirements for 3D cell-laden matrices, low-flow conditions compatible with slow cellular responses, and manufacturing feasibility. Since excessive hydrodynamic force impairs cell viability, a channel depth of 0.5 mm was selected to reduce shear-stress exposure to soft hydrogel-encapsulated cells. Further discussion regarding channel-depth selection is provided in Section S2 of the Supplementary Materials.

### 5.2 Influence of flow rate on velocity and concentration distribution

To evaluate the effect of inlet flow rate on chamber–level concentration gradients under fixed inlet solute concentration, parameter–sweep simulations were conducted across inlet flow rates ranging from 0.1 to 30 μL/min. Velocity and concentration profiles were extracted along the transverse centerline of the culture chamber (Fig. 2b–c).

The transverse velocity profile exhibited a characteristic bimodal distribution: velocity decreased progressively toward the chamber center, reaching its minimum at the geometric midpoint. Peak velocity magnitudes within the chamber scaled monotonically with inlet flow rate. Transverse velocity heterogeneity was pronounced at high flow rates (5–30 μL/min) but became negligible at low flow rates (0.1–2 μL/min).

As shown in Fig. 2d, the transverse concentration gradient along the chamber centerline displayed flow–rate–dependent spatial heterogeneity. Within the proximal region (0.75– 1.55 mm along the transverse centerline), gradient magnitude increased monotonically with rising inlet flow rate (0.1–30 μL/min). In contrast, the distal region (1.55–2.9 mm) exhibited a divergent response: low flow rates (0.1–7.5 μL/min) maintained a positive correlation between flow rate and gradient magnitude, whereas high flow rates (10–30 μL/min) reversed this trend, with gradient magnitude declining as flow rate increased. This bifurcation indicates that elevated flow rates amplify concentration gradients in proximal regions (reflected by steeper concentration profiles per unit length) while suppressing gradients in distal zones.

### 5.3 Effects of channel structure on flow velocity and solute concentration distribution

The proposed multilayer microfluidic chip possesses structural interchangeability, since its flow-channel architecture is independently integrated within the PMMA layer. Accordingly, alternative gradient profiles can be conveniently obtained by simply replacing the patterned PMMA layer with different channel designs. In this study, two typical channel configurations were numerically compared: a conventional curved channel and a channel embedded with Tesla valve structures (Fig. 3). For quantitative comparison, 100 uniformly distributed sampling points were extracted along the central traverse line of each of the four chambers in COMSOL, yielding velocity and concentration profiles for both structural designs.

Consistent with the aforementioned results, both channel structures produced typical bimodal velocity distributions within individual chambers. Overall, the Tesla-valve channel yielded a slightly but consistently lower flow velocity than the conventional curved channel. Statistical quantification showed that the average flow velocity of the Tesla-valve structure was 6.48198×10⁻⁵ m/s, compared with 6.74943×10⁻⁵ m/s for the conventional structure, corresponding to a velocity reduction of approximately 4%.

Solute concentration exhibited a gradual left-to-right decreasing trend across all chambers for both designs. In the first and second chambers, the two channel structures produced nearly identical maximum concentrations at the left chamber regions, whereas the Tesla-valve structure maintained higher residual concentrations at the right low-concentration regions. In contrast, in the third and fourth chambers, comparable maximum concentrations were observed on the right sides, while the Tesla-valve structure generated slightly lower peak concentrations on the left sides.

Quantitative concentration statistics further confirmed that the Tesla-valve architecture barely alters the overall concentration characteristics along the channel centerline. The Tesla-valve channel produced an average concentration of 4.92361×10⁻⁸ mol/m³ with a standard deviation of 3.32915×10⁻⁸ mol/m³, while the conventional curved channel yielded a mean concentration of 4.92595×10⁻⁸ mol/m³ and a standard deviation of 3.29734×10⁻⁸ mol/m³. The two configurations exhibited nearly identical global concentration statistics across the multi-chamber platform, demonstrating that the Tesla-valve units can reduce flow velocity without disrupting the established step-wise concentration-profile.

Tesla-valve structures alter the hydraulic resistance of feeding branches and redirect fluid into vertical bypass passages. At an inlet flow rate of 30 μL/min, the centerline flow velocity shows only a modest (∼4 %) reduction compared with the conventional curved-channel configuration. Simulated concentration profiles (Fig. 3d) show distinct differences between the two designs near the minimum-concentration region within each culture chamber. The conventional curved channel exhibits an abrupt transition, where concentration rapidly plateaus across the right half of the chamber. In contrast, the Tesla-valve-integrated configuration displays a gradual transition toward the concentration plateau on the chamber’s right side.

Multi-flow-rate parametric simulations of the second chamber (Fig. S3) reveal flow-rate-dependent concentration profiles. Increasing inlet flow steepens the central gradient slope, accompanied by stronger nonlinear distortion at both chamber-ends, deteriorated profile symmetry, and a reduced spatial range of nearly linear concentration gradients. The favorable features of the Tesla-valve design, including suppressed intra-chamber advection and gradual concentration transitions, are maintained for inlet flow rates below 10 μL/min but degrade markedly at higher flow rates. At low flow rates of 0.1–0.5 μL/min, broad linear gradient regions with minimal end-region distortion are achieved.

The simulation results demonstrate that the flow velocity and concentration-gradient interval inside the chamber can be tuned by modifying the PMMA channel geometry.

### 5.4 Fabrication of the microfluidic chip and concentration gradient visualization

As shown in Fig. 4, all chip components were fabricated and assembled, and the microfluidic chip was operated using an injection micropump. Two 1 mL syringes were first loaded with blue and red colored water, respectively. Silicone tubing was then attached to the syringe needles, and the syringes were mounted on the injection micropump. The opposite end of each silicone tubing was connected to a stainless-steel capillary at the inlet of the microfluidic channel. The total injection volume was set to 60 μL at a flow rate of 5 μL/min. The colored water served dual purposes: it visualized the concentration gradient and simultaneously verified the absence of leakage at all connections. For constructing the hydrogel stiffness interface within the microfluidic chip, a 3D bioprinter was used to deposit Pluronic F127 hydrogel encapsulating the internal dextran hydrogel. The resulting interface between the two hydrogel types within the microfluidic chamber is shown in Fig. 5.

**Fig. 4.**
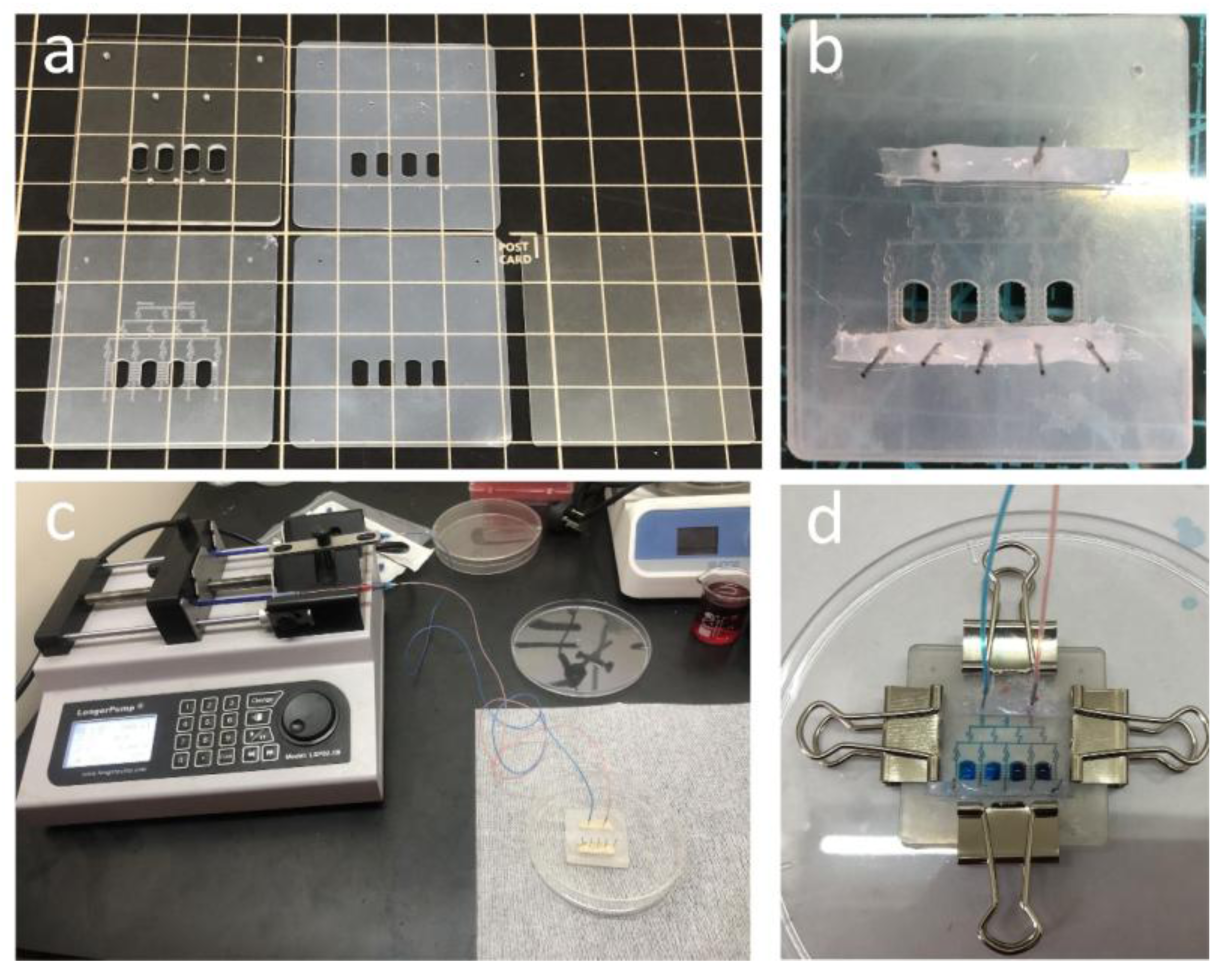
Chip assembly and fluid leakage test. (a) Individual layered components of the microfluidic chip; (b) Assembled microfluidic chip; (c) Experimental setup for fluid delivery; (d) Visualization of colored fluid within the chip.

**Fig. 5.**
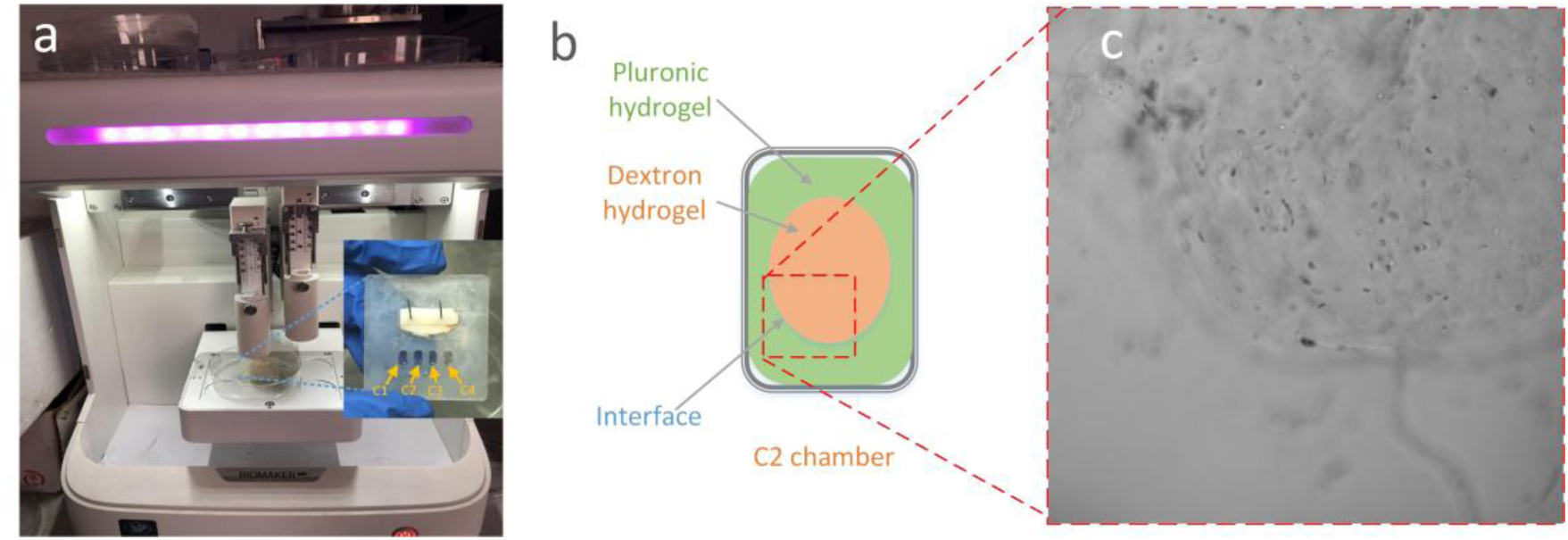
Hydrogel interface fabricated inside microfluidic chambers. (a) 3D bioprinter depositing Pluronic F127 hydrogel to encapsulate internal dextran hydrogel; (b) Schematic and micrograph of the resulting hydrogel interface formed by this bioprinting strategy.

### 5.5 Cell viability in microfluidic chip

The microfluidic chip developed herein was designed to generate solute concentration gradients and investigate their effects on cellular behavior. SKOV3 cells were first encapsulated within 3D dextran hydrogel and then loaded into the culture chambers of the microfluidic device. A syringe was filled with 1 mL of RPMI-1640 complete medium and connected to a programmable syringe pump for perfusion. After 24 h of on-chip culture, cells in each chamber were stained for live/dead assays. As shown in Fig. 6, green fluorescence denotes live cells, while red fluorescence indicates dead cells. Following 24 h of culture of 3D-encapsulated SKOV3 cells inside the chip chambers, cell viability remained high across all chambers, with only a small fraction of dead cells observed. The time-dependent cellular growth profile within representative chamber C2 is presented in Fig. 7.

**Fig. 6.**
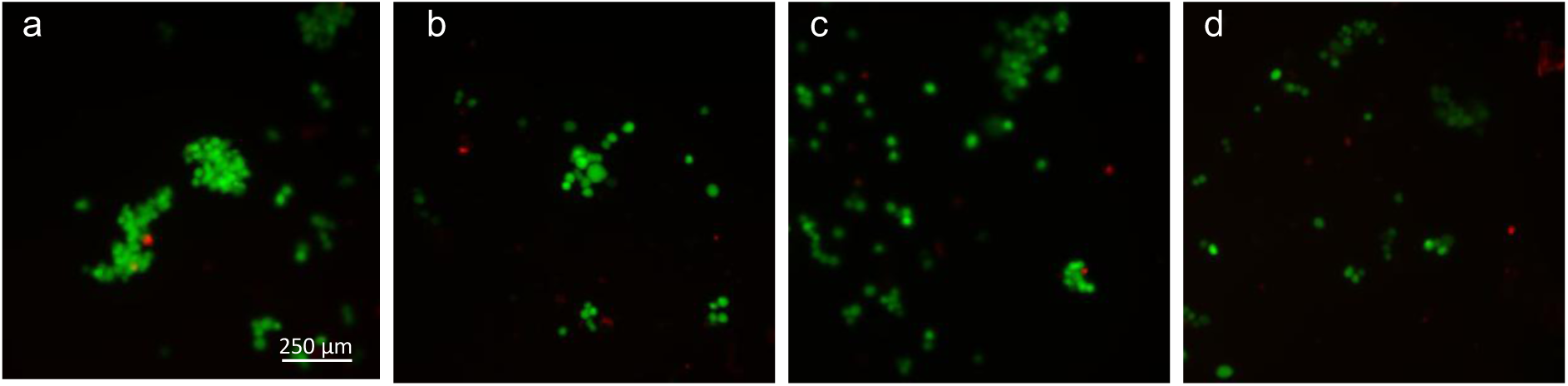
Fluorescence micrographs of live/dead cell staining obtained from the four culture chambers.

**Fig. 7.**
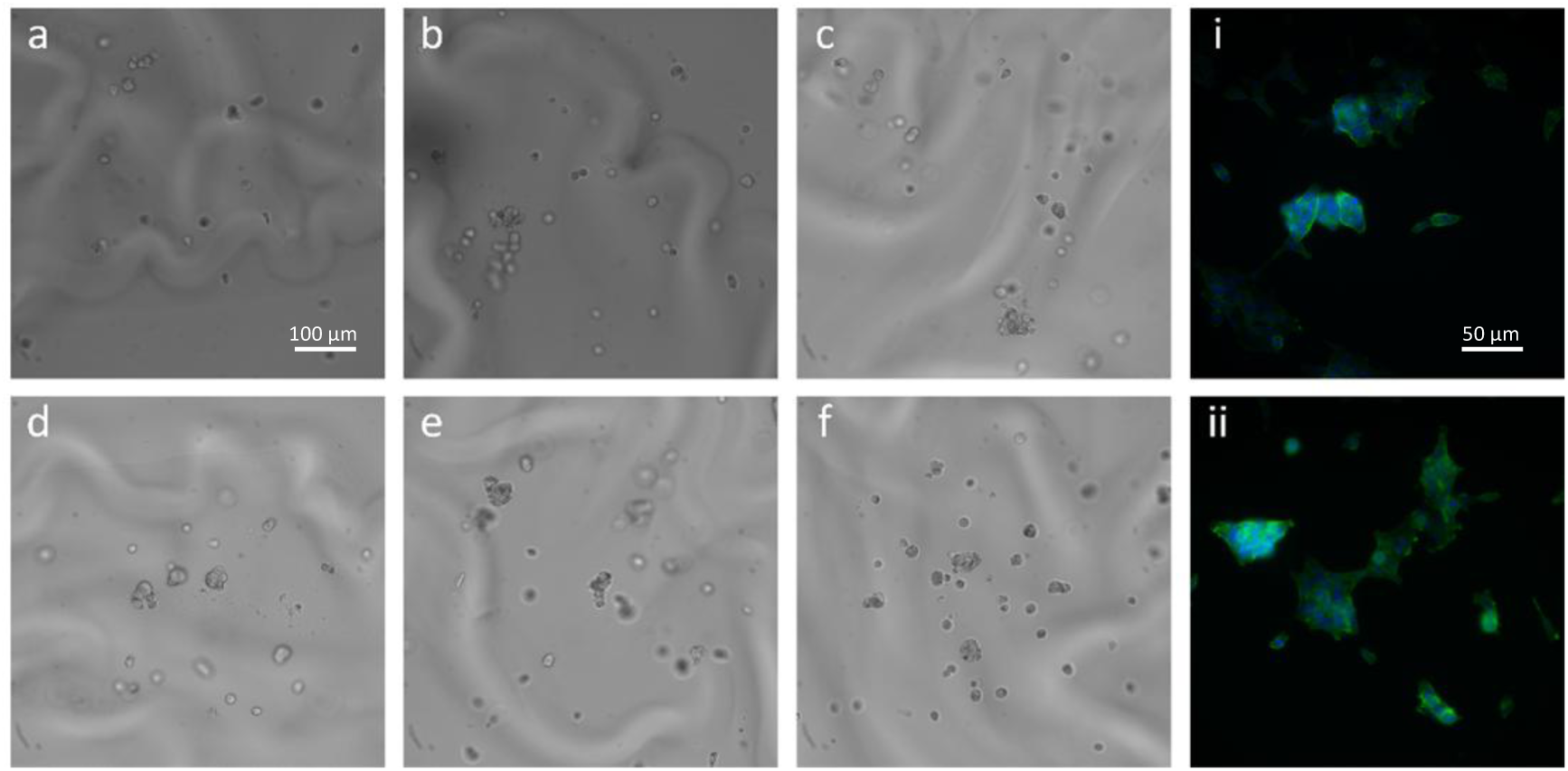
Bright-field and fluorescence micrographs of hydrogel-embedded SKOV3 cells in chamber C2. (a–c) 4 h, 12 h and 24 h culture for cells in the conventional curved-channel configuration; (d–f) 4 h, 12 h and 24 h culture for cells in the Tesla-valve channel configuration. (i, ii) Corresponding magnified fluorescence views at 24 h. Green: F-actin; blue: DAPI.

### 5.6 Effect of TGF-β1 concentration on the growth status of SKOV3 cells

The microfluidic chip produces distinct TGF-β1 concentration distributions in each chamber. SKOV3 cells were cultured for 48 h in two microenvironments: an on-chip experimental group and a 24-well plate control group using conventional dextran hydrogel with 10 ng/mL TGF-β1.

Bright-field micrographs of cells within the 3D hydrogels at 12 h, 24 h and 48 h are displayed in Fig. 8. Micrographs of the on-chip group were acquired from the leftmost chamber of the device. After 12 h of culture, cells in the chip chamber had not yet extended pseudopodia, while visible pseudopodial protrusion was observed at 48 h. This observation suggests cells required an extended period to locate adhesion sites within the confined, tailored microenvironment of the chip chamber. In contrast, cells in the hydrogel samples cultured in well plates exhibited partial elongation. Over the 48 h incubation period, most cells in the well-plate group developed pseudopodia with minimal cell aggregation, whereas cells inside the chip chambers displayed prominent aggregation from 24 h to 48 h.

**Fig. 8.**
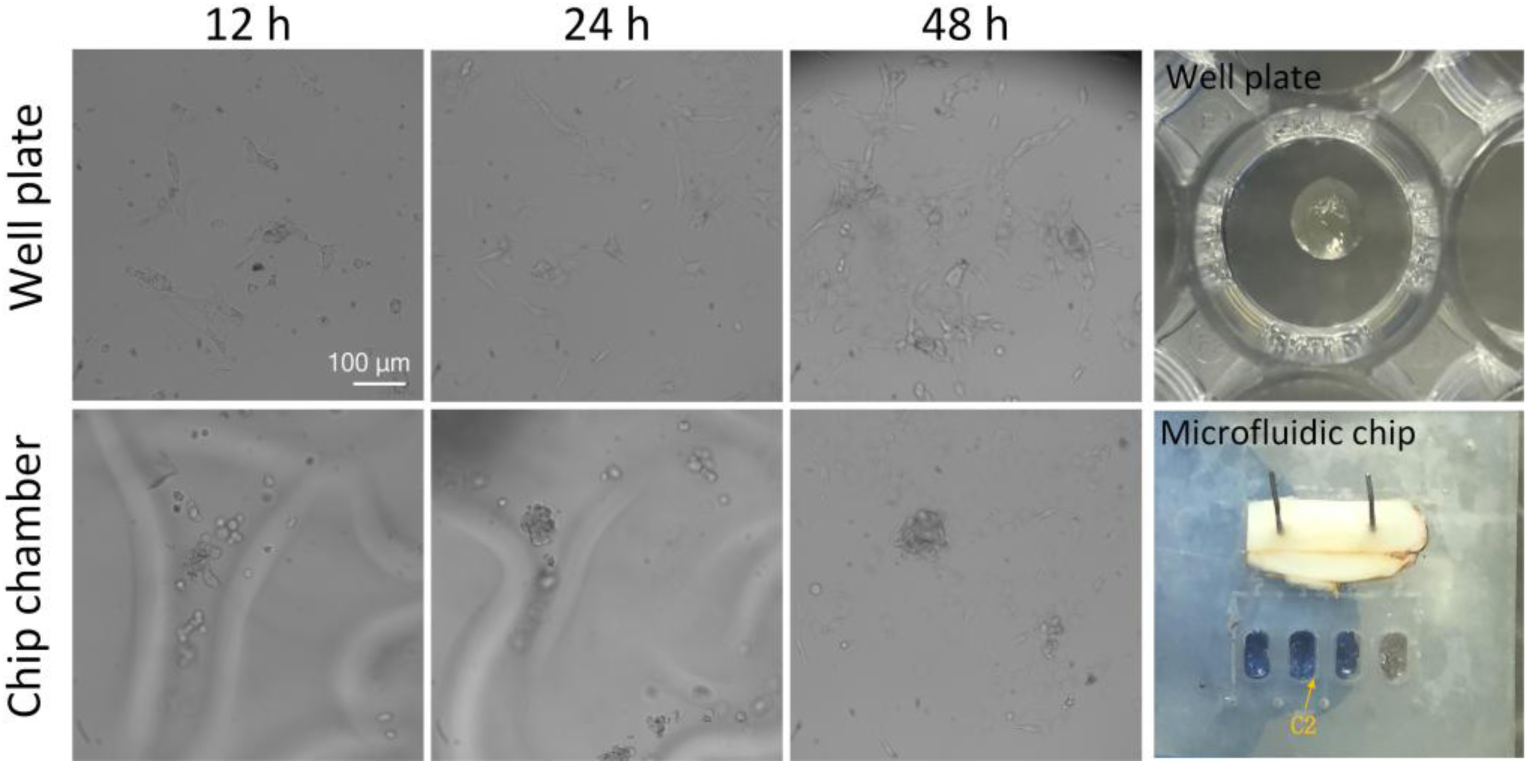
Morphology of SKOV3 cells at different time points under TGF-β1 stimulation.

After 48 h of incubation, gradual degradation of the dextran hydrogel was observed. This degradation was driven by matrix metalloproteinases (MMPs) secreted by SKOV3 cells, which specifically cleave MMP-sensitive peptide sequences within the cross-linking CD-linkers. Cleavage of CD-linkers disrupts the hydrogel network and impairs local mechanical integrity, allowing SKOV3 cells to migrate within the matrix. Meanwhile, progressive hydrogel erosion caused partial detachment of encapsulated cells. Some cells sedimented onto the bottom culture surface and proliferated either within residual hydrogel fragments or on the thin gel layer attached to the substrate.

Within the microfluidic chip, the TGF-β1 concentration decreases sequentially from the leftmost to the rightmost chamber. The TGF-β1 concentration ranges in chambers C1, C2, C3 and C4 are 8–10 ng/mL, 5–8 ng/mL, 2–5 ng/mL and 0–2 ng/mL, respectively. The growth phenotypes of SKOV3 cells cultured for 48 h inside the four chip chambers are presented in Fig. 9. The size and abundance of cell aggregates in chambers C1, C2 and C3 were greater than those observed in chamber C4, indicating that cells tended to form spherical aggregates upon stimulation by relatively high TGF-β1 concentrations (approximately 5–10 ng/mL). In this hydrogel system, high TGF-β1 concentrations promoted the aggregation of SKOV3 cells, with relatively few cells migrating and spreading into the surrounding matrix. In contrast, lower TGF-β1 concentrations suppressed aggregate formation and facilitated cell spreading and migration.

**Fig. 9.**
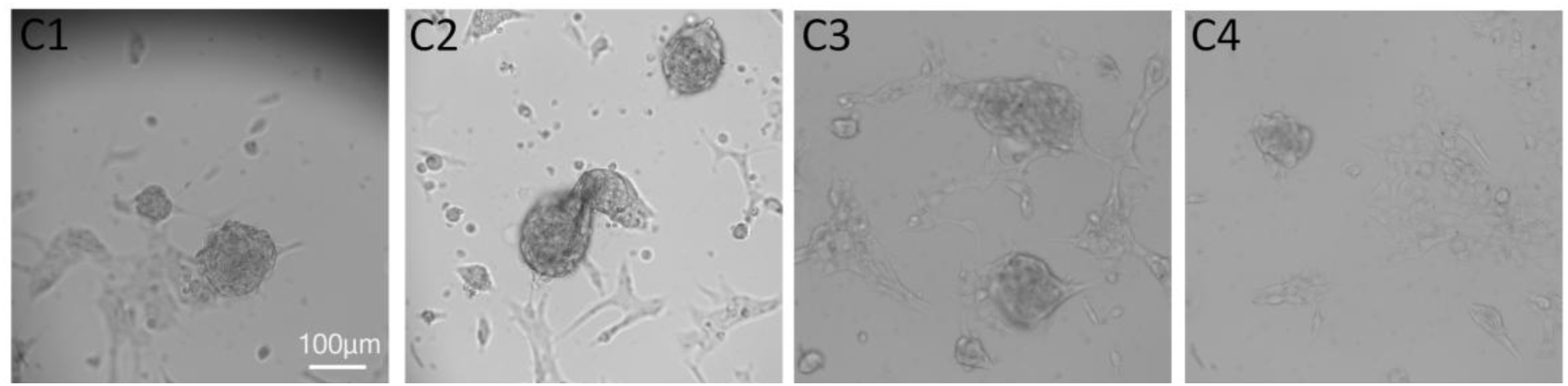
Morphology of SKOV3 cells in each microfluidic chamber after 48 h of TGF-β1 stimulation.

### 5.7 Effect of TGF-β1 concentration on adhesion protein and cytoskeletal protein expression in SKOV3 cells

SKOV3 cells cultured for 48 h in 24-well plates and microfluidic chip chambers were fixed and stained for the cytoskeletal protein F-actin and adhesion protein N-cadherin, as shown in Figs. 10 and 11. As illustrated in Fig. 10, most SKOV3 cells adhered to the bottom of the hydrogel layer in well-plate samples. Prominent lamellipodia observed in F-actin staining indicated active cell migration. By contrast, F-actin fluorescence revealed that cells inside microfluidic chip chambers (Fig. 11) mainly formed spherical or filamentous aggregates with obvious spherical clusters. Although F-actin expression remained detectable in these cells, they exhibited fewer lamellipodia compared with cells cultured in well plates. Overall, cells within microfluidic chambers underwent more extensive aggregation relative to well-plate cultures. This phenotype may originate from the confined microenvironment within the chip: limited gas permeability (O₂, CO₂) and reduced access to culture medium per unit volume of hydrogel matrix. Furthermore, the graded TGF-β1 concentration also contributes to this cellular phenotype. Consistent with this trend, cell aggregates were less abundant in chambers C3 and C4 than in C1 and C2.

**Fig. 10.**
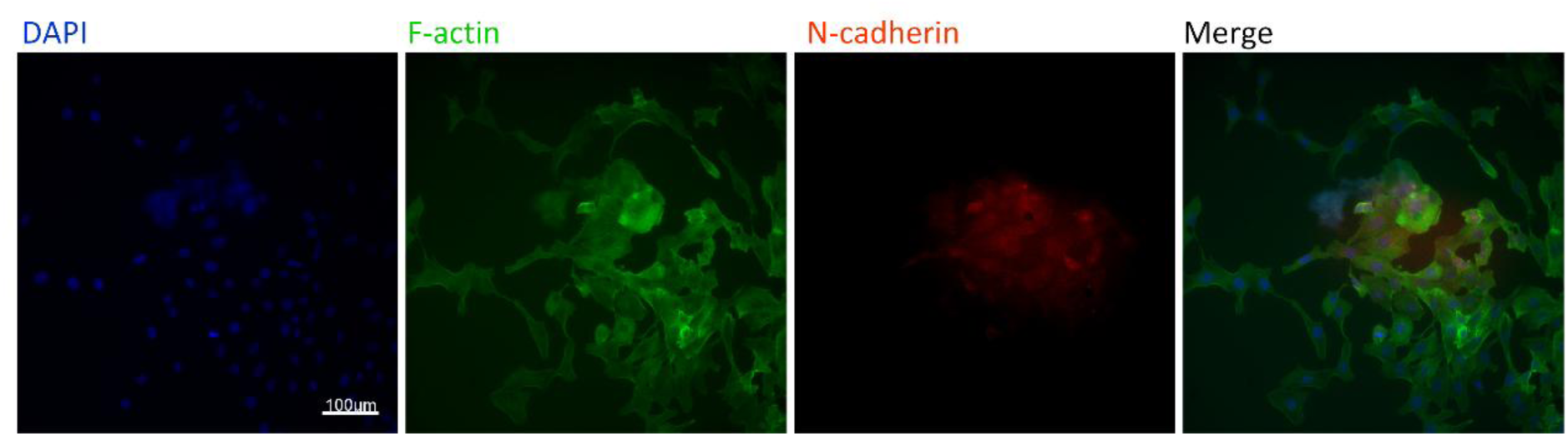
**Fluorescence micrographs of target protein expression in hydrogel-embedded SKOV3 cells after 48 h of TGF-β1 stimulation in well-plate cultures.**

**Fig. 11.**
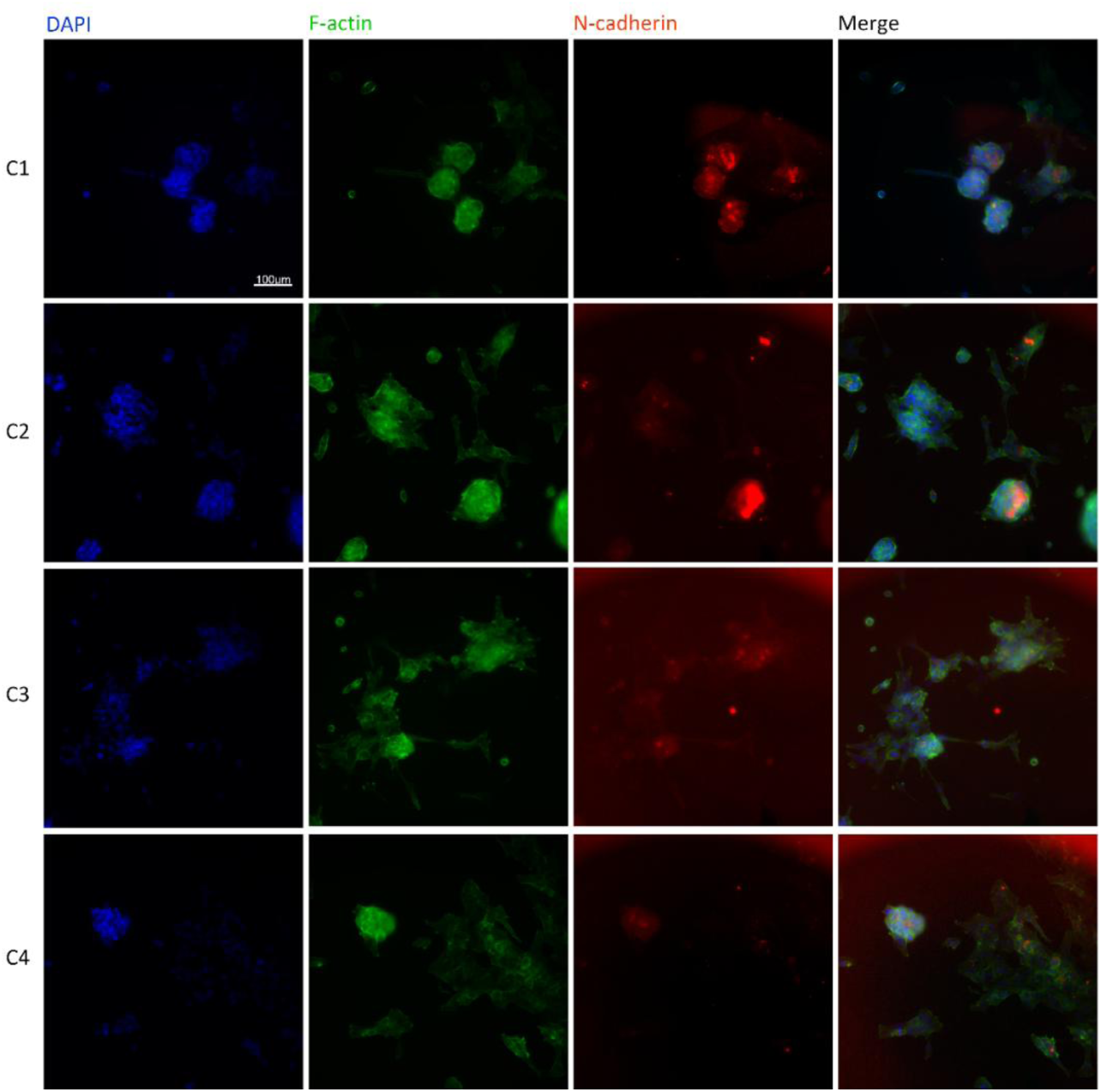
Fluorescence micrographs of stained SKOV3 cells embedded in hydrogels within microfluidic chips after 48 h of TGF-β1 stimulation.

Figures 11 and 12 present fluorescence staining of hydrogel-embedded SKOV3 cells following 48 h of TGF-β1 stimulation. In homogeneous dextran hydrogels, SKOV3 cells exhibited spontaneous spreading and migratory behavior, accompanied by abundant F-actin filament formation. Two typical cell aggregation morphologies were observed, including spherical clusters and elongated filament-like clusters. After the fabrication of Pluronic-based hydrogel stiffness interfaces via 3D bioprinting, SKOV3 cells tended to migrate perpendicularly toward the mechanical boundary through durotaxis. Filopodia were actively formed during early migratory processes. However, most cells eventually accumulated adjacent to the stiffness interface after 48 h and showed no further distant migration. Cells residing at the stiffness boundary predominantly exhibited aggregated morphology with reduced filopodial protrusions. Under this mechanical boundary condition, only spherical cell clusters with detectable F-actin expression were observed (Fig. 12).

**Fig. 12.**
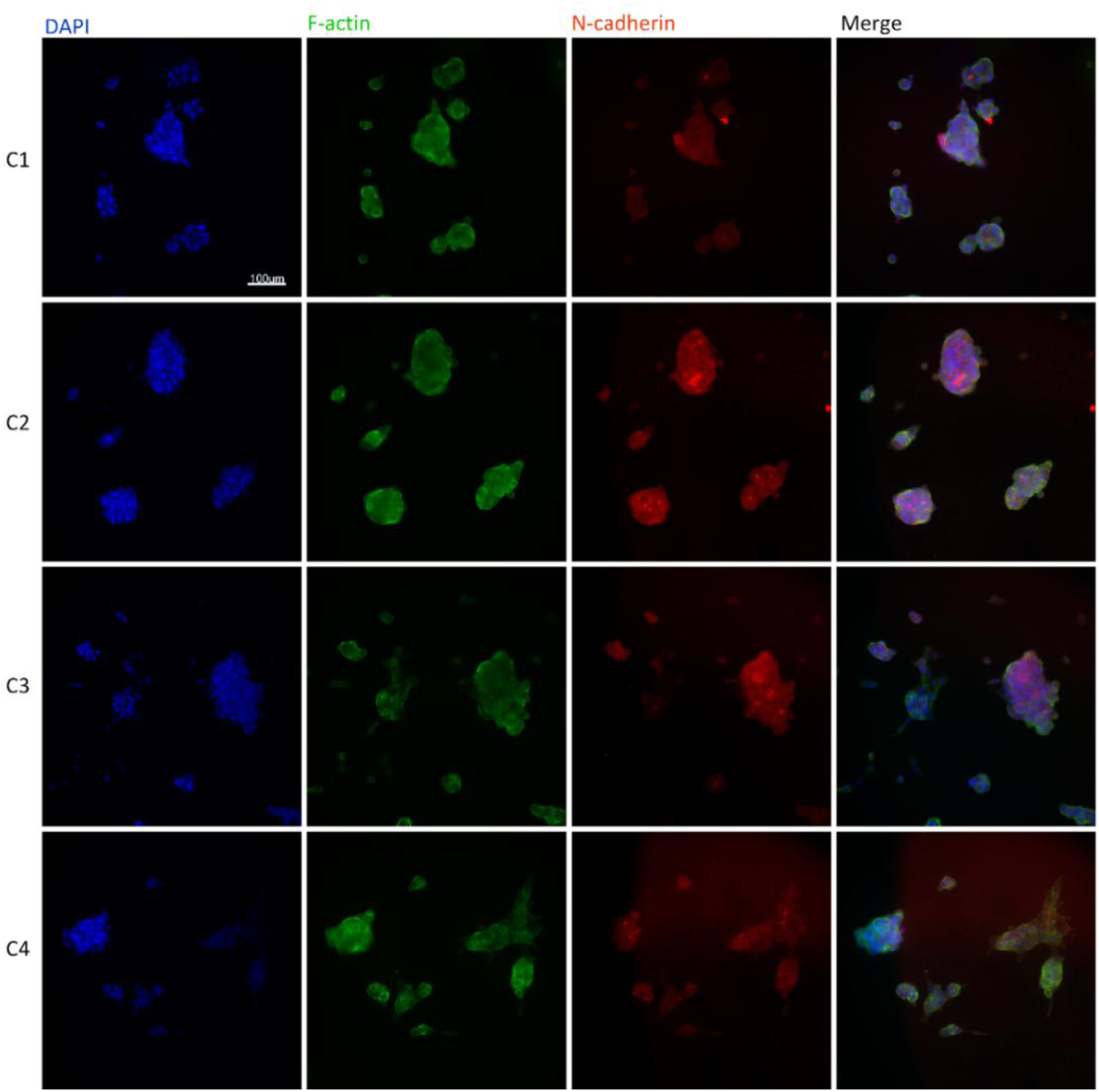
Fluorescence micrographs of stained SKOV3 cells at the hydrogel stiffness interface within microfluidic chips after 48 h of TGF-β1 stimulation.

N-cadherin serves as a canonical mesenchymal biomarker closely associated with the epithelial-to-mesenchymal transition (EMT) process. As quantified in Fig. 14, N-cadherin exhibited an expression pattern analogous to that of F-actin. Within the four microfluidic culture chambers, N-cadherin fluorescence intensity decreased progressively from left to right, and higher fluorescence signals were observed in the stiffness-interface groups relative to the homogeneous intra-chip gel groups. For the first chamber with the highest TGF-β1 concentration, the F-actin fluorescence intensity of SKOV3 cells in the homogeneous-gel group C1 was comparable to that in the well-plate control group, and both values were slightly lower than those of the stiffness-interface group C11. Notably, groups at the stiffness-interface displayed smaller standard deviations for N-cadherin fluorescence intensity, indicating more stable N-cadherin expression among SKOV3 cells under this microenvironmental condition.

## 6. Discussion

### 6.1 Mechanisms of mass transport modulation by Tesla-valve integrated microfluidic architecture

Tesla-valve structures raise the hydraulic resistance of each feeding branch and increase pressure on both the high-concentration (left-hand) and low-concentration (right-hand) sides of each culture chamber. At a fixed total inlet flow rate, more fluid is diverted into vertical bypass passages instead of crossing the chamber’s central region through horizontal narrow connecting channels. This leads to only a modest (∼4 %) reduction in centerline flow velocity and moderately suppresses horizontal advection, making molecular diffusion dominate solute transport. This effect yields a gentler concentration-profile slope and stabilizes the concentration gradient, which is consistent with the simulation observation that Tesla-valve units generate more gradual concentration-gradient intervals. This smoother transition avoids sharp concentration inflection points, which is biologically favorable for cell culture by mitigating abrupt chemical shifts to cellular microenvironments.

The above-mentioned simulations (Fig. 3) were carried out at 30 μL/min, a relatively high inlet flow rate. Although the performance of the Tesla-valve-integrated design is partially compromised under this condition, the observable contrast between the two channel layouts is sufficiently pronounced to clarify the underlying transport mechanisms. Notably, stable concentration gradients can still be formed across all four culture chambers even at this high-flow-rate condition, which demonstrates that the Tesla-valve-equipped configuration can effectively suppress excessive intra-chamber convection. It can be inferred that even smoother and more stable concentration profiles will be achieved under reduced-flow-rate regimes where molecular diffusion dominates mass transport.

Nevertheless, practical cell-culture experiments require far lower flow magnitudes. Strong intra-chamber convection should be avoided for 3D hydrogel-embedded cell assays: convective disturbance introduces undesired fluid-shear mechanical cues that may interfere with cellular mechanotransduction, distorts the pre-designed chemical gradient field, and washes away locally accumulated autocrine-paracrine signaling molecules. All these artifacts would confound the biological readouts of cell migration and drug response. Moreover, cellular responses to biochemical stimuli occur on a slow-timescale of tens of hours, calling for a quasi-static, diffusion-dominated microenvironment. Accordingly, an inlet flow rate below 1 μL/min was selected for the following biological cell-culture experiments to balance stable gradient performance and practical chip operation.

### 6.2 Biological responses of SKOV3 cells under TGF-β1 concentration gradient

F-actin is a major cytoskeletal protein whose polymerization and depolymerization are tightly regulated by actin-binding proteins (ABPs)^[18]^. Active ABP-mediated F-actin polymerization occurs at the leading edge of migrating cells, and the corresponding fluorescence intensity was quantitatively analyzed in this study (Fig. 13). N-cadherin was selected as the primary mesenchymal biomarker to evaluate EMT progression, while F-actin staining was used to characterize cytoskeletal remodelling, a downstream cellular phenotype accompanying EMT. The formation of lamellipodia and filopodia reflects cytoskeletal remodelling and facilitates EMT progression, which can enhance the metastatic potential of cancer cells^[19]^. Nevertheless, these protrusions alone cannot confirm EMT activation; upregulation of mesenchymal markers such as N-cadherin is required to verify EMT progression.

**Fig. 13.**
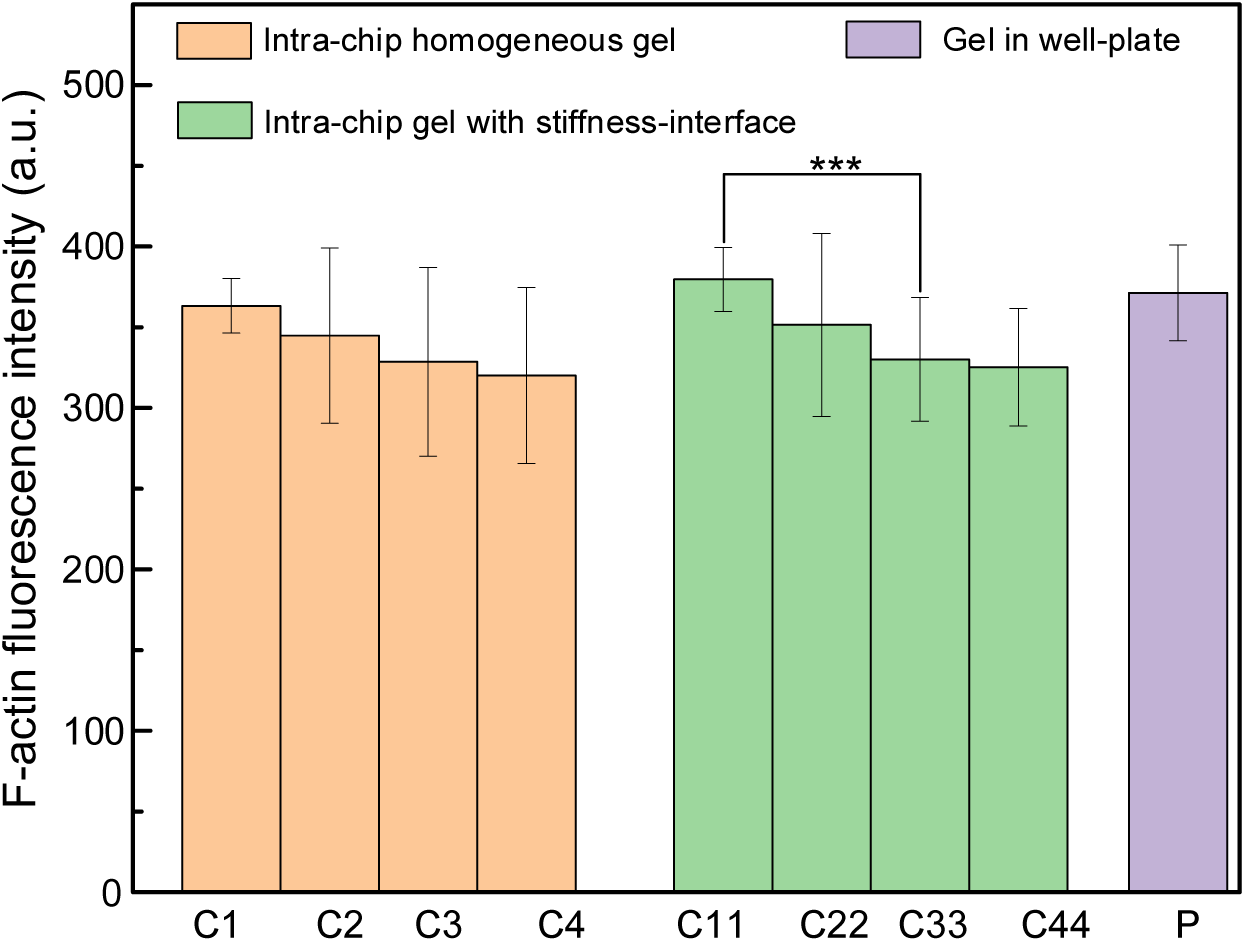
Quantification of F-actin fluorescence intensity (a.u., mean gray values obtained from ImageJ analysis) for SKOV3 cells embedded in hydrogels after 48 h treatment with TGF-β1. Intra-chip homogeneous gel (C1-C4): cells within uniform hydrogel regions of the microfluidic chip. Stiffness-modulated gel interface (C11-C44): cells located at the hydrogel stiffness interface inside the chip. Gel in well-plate (P): hydrogel constructs incubated directly in well-plates without the microfluidic device. Groups C1 and C11 correspond to the first chamber with the highest average TGF-β1 concentration. C1-C4 and C11-C44 represent varied chemical gradient conditions. *** *p* < 0.007, *n* = 20. Error bars represent mean ± SD.

**Fig. 14.**
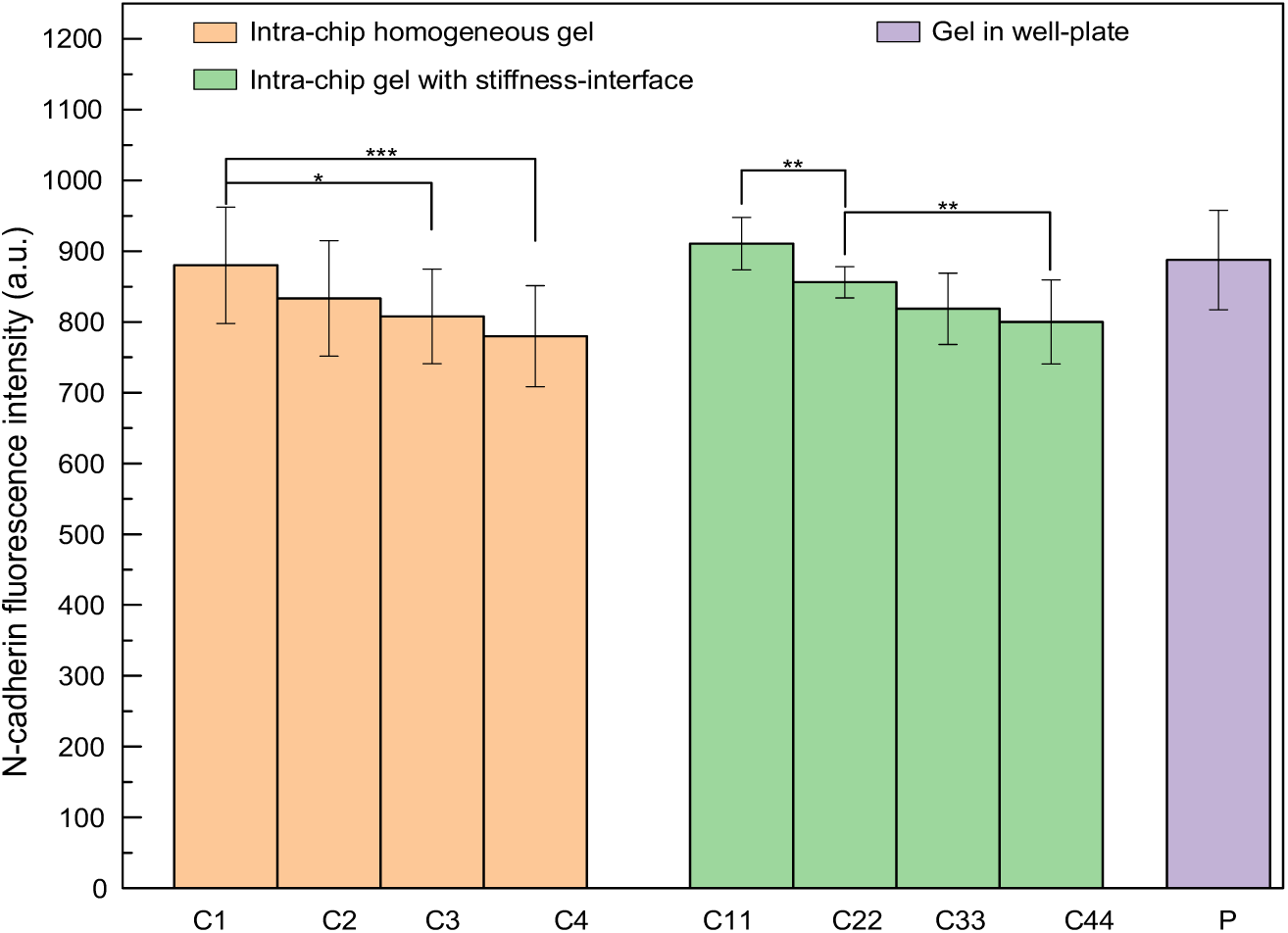
Quantification of N-cadherin fluorescence intensity (a.u., mean gray values obtained from ImageJ analysis) for SKOV3 cells embedded in hydrogels after 48 h treatment with TGF-β1. Intra-chip homogeneous gel (C1-C4): cells within uniform hydrogel regions of the microfluidic chip. Intra-chip gel with stiffness-interface (C11-C44): cells located at the hydrogel stiffness interface inside the chip. Gel in well-plate (P): hydrogel constructs incubated directly in well-plates without the microfluidic device. Groups C1 and C11 correspond to the first chamber with the highest average TGF-β1 concentration. * *p* < 0.05 (C1 vs C3), ** *p* < 0.01 (C11 vs C22, C22 vs C44), *** *p* < 0.001 (C1 vs C4), *n* = 20. Error bars: mean ± SD.

Within the microfluidic gradient chip, F-actin expression decreased gradually from the high-TGF-β1 chamber (C1) to the low-TGF-β1 chamber (C4). Moreover, cells located at the hydrogel stiffness interface exhibited significantly higher F-actin levels than those in homogeneous hydrogel regions under identical biochemical conditions. In the chamber with the highest TGF-β1 concentration, the F-actin intensity of cells in the homogeneous gel group (C1) was slightly lower than that in the stiffness-interface group (C11) and the well-plate control. Group C11 showed a cytoskeletal activation level comparable to that of the conventional 3D culture group. In line with this morphological signature, SKOV3 cells cultured in homogeneous intra-chip hydrogels developed obvious lamellipodia, enabled by abundant RGD adhesion sites within the hydrogel network that support cell spreading and migration.

TGF-β1 is a key upstream regulator that induces EMT and malignant progression in ovarian cancer via the canonical Smad signaling pathway^[20]^. Specifically, TGF-β1 stimulation activates Smad2/3 signaling, upregulates mesenchymal markers including N-cadherin and vimentin, and suppresses epithelial markers such as E-cadherin, thereby driving cytoskeletal remodeling and enhancing cell invasive capacity^[21, 22]^. Consistent with these reported mechanisms, our experimental results demonstrated a clear TGF-β1 concentration-dependent EMT response. Higher TGF-β1 levels corresponded to elevated N-cadherin expression and promoted the formation of compact spherical cell aggregates, a phenotypic hallmark of aggressively invasive ovarian cancer cell populations^[23]^. Although the gradient-induced expression variation across chambers was moderate, the progressive reduction of F-actin and N-cadherin from C1 to C4 strongly confirms that decreasing TGF-β1 stimuli gradually weaken EMT activation and cytoskeletal remodeling.

As quantified in Fig. 13 and Fig. 14, F-actin and N-cadherin fluorescence intensities of SKOV3 cells declined progressively from chamber C1 to C4 and from C11 to C44, matching the decreasing TGF-β1 concentrations across the chemical gradient. This observation demonstrates that TGF-β1 drives cytoskeletal remodelling and EMT-related molecular reprogramming in a concentration-dependent manner within 3D hydrogel matrices. Higher TGF-β1 concentrations promoted F-actin assembly and elevated N-cadherin levels, favouring the acquisition of a mesenchymal, pro-invasive phenotype.

Notably, under equivalent TGF-β1 concentrations, cells residing at the stiffness-modulated interface exhibited higher F-actin and N-cadherin signals than those cultured in intra-chip homogeneous hydrogels, as exemplified by the comparison between C11 and C1. These results indicate synergistic crosstalk between biochemical stimulation (TGF-β1 gradient) and mechanical microenvironmental cues (stiffness-interface-mediated durotaxis). Although cells accumulated adjacent to the stiffness boundary and failed to migrate further away, they retained elevated expression of migration- and EMT-associated markers. This implies that cancer cells may sustain an invasive molecular signature even when physical migration is mechanically constrained. Moreover, comparable F-actin levels between group C11 and the off-chip gel-in-well-plate control (P) confirm that the confined microfluidic chamber itself does not intrinsically suppress cytoskeletal activation or EMT responses.

### 6.3 Biological significance of the microfluidic platform

In vivo tumor tissues feature both spatial gradients of soluble cytokines and mechanical heterogeneity of the extracellular matrix. Our findings highlight that biochemical and mechanical signals jointly shape the malignant phenotype of ovarian cancer cells. In our 3D in vitro model, cells located at matrix stiffness boundaries retained elevated expression of EMT-associated markers despite limited long-range migration, suggesting that mechanical boundary cues may support the formation of primed, metastasis-competent cell populations within the primary tumor microenvironment. Conventional well-plate assays usually deliver uniform cytokine doses and cannot recapitulate combined biochemical gradients and defined mechanical interfaces simultaneously. Benefiting from the reconfigurable architecture of our microfluidic chip, coupled biochemical-mechanical tumor-microenvironment cues can be reconstructed in vitro, offering a useful platform for dissecting the multifactorial regulation of ovarian cancer cell malignant behavior.

## 6. Conclusion

In this work, we designed and fabricated a reconfigurable multilayer microfluidic chip comprising a top PMMA cover, two silicone sealing membranes, a modular PMMA channel layer, and a bottom glass substrate. By altering channel layouts and chamber combinations, this PMMA structure enables tunable flow fields and biomolecule concentration gradients. COMSOL Multiphysics finite element simulations were used to characterize creeping flow and solute transport inside the porous hydrogel microenvironment. Simulations identified 0.5 mm as the optimal channel depth for generating low-velocity flow fields and stable, gradual concentration gradients; increasing inlet flow rate steepened the spatial gradient within culture chambers.

After numerical characterization, the microfluidic device was assembled and experimentally validated. Functional tests confirmed reliable gradient formation and high viability of 3D hydrogel-encapsulated SKOV3 cells cultured on-chip. Using this platform, we demonstrated that the TGF-β1 concentration gradient regulates cytoskeletal remodeling and the expression of the mesenchymal marker N-cadherin in SKOV3 cells. Elevated EMT-associated molecular signatures were detected under higher TGF-β1 concentrations and at hydrogel matrix stiffness interfaces.

This microfluidic platform is easy to assemble, low-cost and highly integrated, with interchangeable structural components for concentration gradient generation. Its capability for 3D cell culture was verified in the present study. The modular PMMA architecture shortens chip fabrication and iteration cycles relative to conventional PDMS microfluidic devices and can be adapted for other biochemical stimuli and cell types in future work. This reconfigurable microfluidic strategy offers a feasible tool to probe cellular behaviors and protein expression under precisely controlled chemomechanical microenvironments.

## Acknowledgments

This work was supported by the National Natural Science Foundation of China (Grant No. 51875170) and the University-Industry Collaboration Project (Grant No. CZ824105716). The authors thank Mr. Erxuan Xiong in our laboratory for assistance with preliminary experimental tests.

