## Supplementary Materials for "Multilayer-assembled microfluidic chip with Tesla-valve channels: Combined biochemical gradient and mechanical stiffness interface for 3D SKOV3 cell culture"

S1 Schematic diagram for each layer of the multi-layer microfluidic chip

The multi-layer microfluidic chip consists of five stacked components, as illustrated in Fig. S1. Layer 1 is the top PMMA cover plate, which contains positioning holes, two inlets (Inlet A and Inlet B) and multiple outlets for fluid delivery and collection (Fig. S1a). Layer 2 is the first silicone sealing film with reserved openings aligned to the culture chambers (Fig. S1b). Layer 3 is the patterned PMMA flow-channel layer, embedding Tesla valve structures, interconnected microchannels and four sequential culture chambers to generate controllable concentration gradients (Fig. S1c). Layer 4 is the second silicone sealing film, which isolates the channel layer from the substrate while retaining the chamber regions (Fig. S1d). Layer 5 is the flat optical glass substrate serving as the bottom support (Fig. S1e). All layers are precisely aligned through positioning holes (also referred to as location holes) and assembled to form the complete microfluidic device (Fig. S1f). The interchangeable design of Layer 3 allows rapid replacement of flow-channel geometries, such as conventional curved channels or Tesla-valve integrated layouts, to tune flow fields and solute concentration profiles inside the culture chambers.

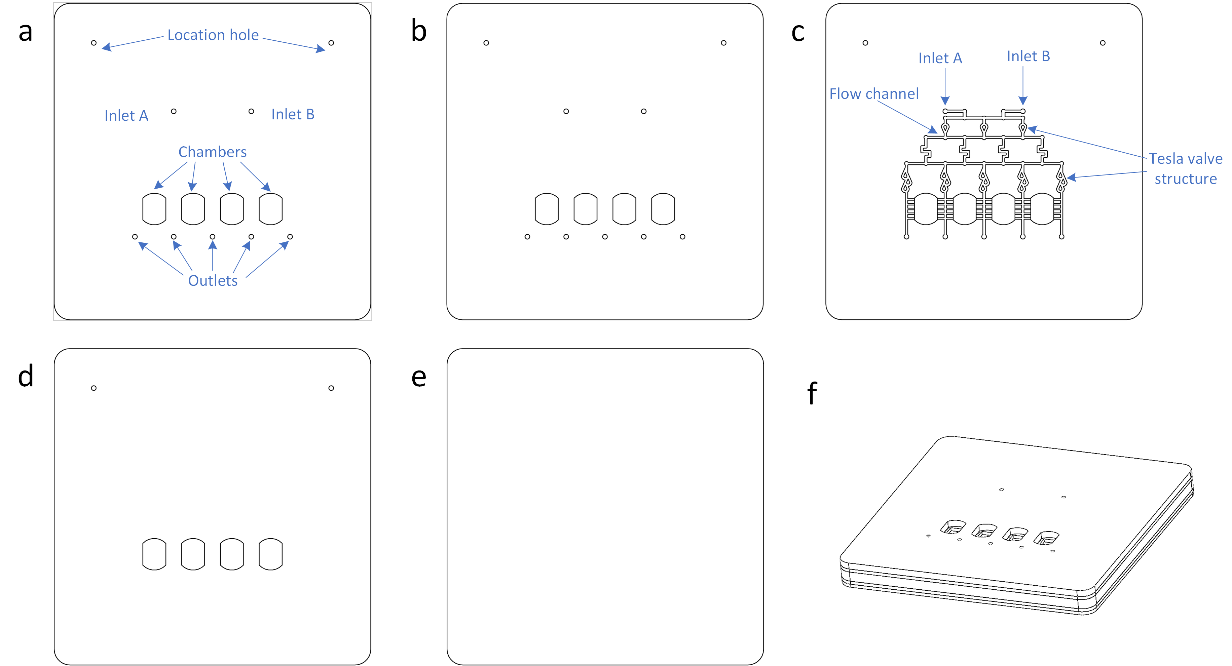

**Fig. S1** Schematic diagram of the multi-layer microfluidic chip. (a) Top PMMA cover layer (Layer 1) with positioning holes, inlets and outlets; (b) The first silicone sealing film (Layer 2); (c) PMMA flow-channel layer (Layer 3) patterned with microchannels, Tesla valve structures and four culture chambers; (d) The second silicone sealing film (Layer 4); (e) Bottom optical glass substrate (Layer 5); (f) 3D schematic view of the fully assembled microfluidic chip.

S2 Simulation for different depths of flow channels and chambers

To optimize the depth of the flow channel/chamber shown in Fig. S2(a), PMMA channels with depths of 0.3 mm, 0.4 mm, and 0.5 mm were evaluated. When the inlet flow rate was set to 30 μL/min, and the substance concentration c was defined as 1×10⁻⁷ mol/m³ and the inlet flow rate is unchanged, the flow rate in the interior of the chamber shows a decreasing trend with the increase of the depth of the PMMA flow channel as shown in Fig. S2(b). When the inlet flow rate is 30 µL/min and the flow channel groove depth is set to 0.5 mm, the internal flow velocity within the chamber is around 0.06mm/s, which is significantly lower than that with groove depths of 0.3 mm and 0.4 mm. In addition, the flow velocities at both ends of the chamber are higher than in the central region, and the case of 0.3/0.4-mm chamber depths create elevated shear stress compared to that of 0.5-mm chamber depths. As shown in Fig. S2(c), the substance concentrations in the chamber shows decreasing trend from left to right, and the concentration distribution over the positions of 1.0~2.0 mm on the centerline of the chamber for the 0.5 mm-depth channel is similar to that of 0.3 mm-depth and 0.4 mm-depth. The concentration gradient produced by the 0.5 mm-depth channel is relatively gentle in the lateral regions (left and right side-areas), whereas it becomes moderately steep in the central region, thereby facilitating effective stimulation of cells within the gel sample located in this central area.

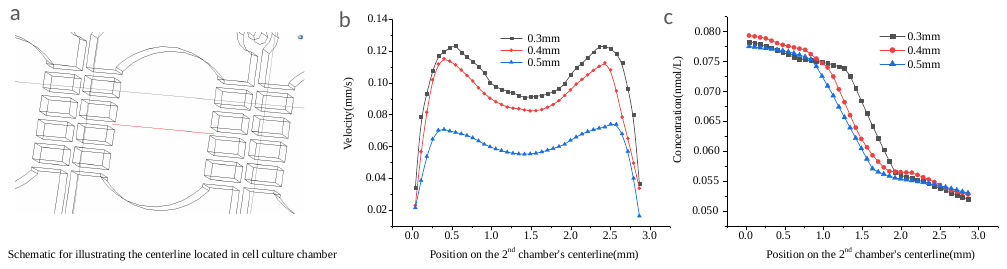

**Fig. S2** Influence of the depth of flow channels (DoF) in the layer 3 on the flow velocity and concentration of stimulating substance. (**a)** schematic for illustrating the centerline of cell culture chamber; **(b)** velocity of centerline in the 2^nd^ chamber with different DoFs; **(c)** substance concentration along the centerline in the 2^nd^ chamber.

**S3 Comparison of valve‑equipped structure and conventional channel structure under different flow‑rate conditions**

To investigate how different flow channel architectures, affect flow and concentration fields within a single reaction chamber, two distinct channel configurations—Tesla valve channels and conventional curved channels—were fabricated by modifying the geometry of the third channel layer. Three volumetric flow rates (1, 10, and 30 μL/min) were implemented to represent low, moderate, and high flow conditions, respectively. Numerical simulations were performed focusing on the second chamber of the microfluidic device.

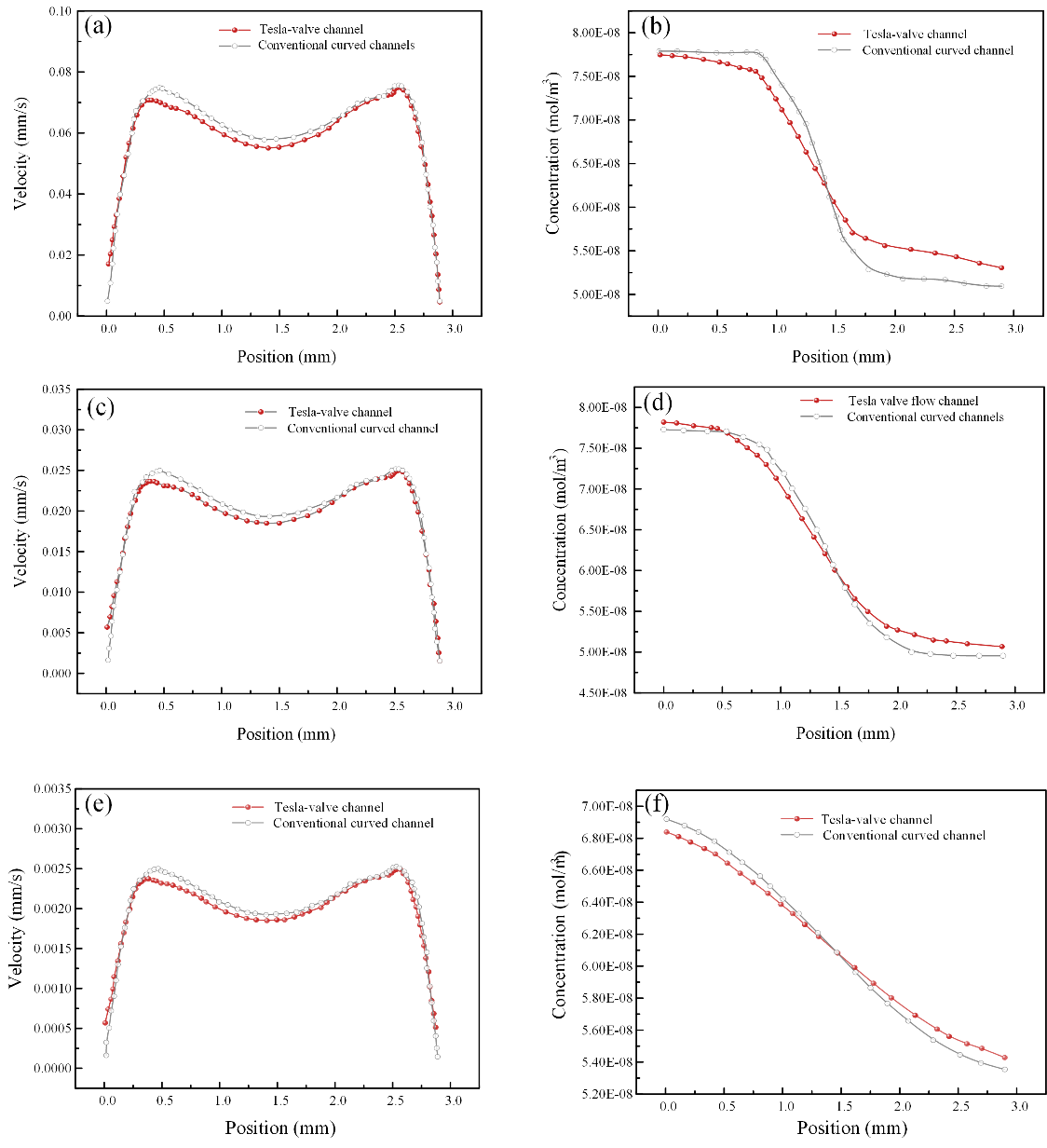

**Fig. S3** Velocity and concentration distribution differences between Tesla-valve and conventional curved flow channels under three inlet flow rates. (a–b) High flow rate of 30 μL/min; (c–d) Moderate flow rate of 10 μL/min; (e–f) Low flow rate of 1 μL/min.

Figure S3 compares velocity and concentration distributions across the two channel designs under varied flow rates. As illustrated in Figs. S3a, S3c and S3e, both the Tesla-valve and conventional curved channels produce double-peak velocity profiles within the second chamber at all tested inlet flow rates.

For the conventional curved channel, the horizontal narrow connecting channels on both sides of the chamber follow a symmetric layout. The flow path length and hydraulic resistance from the chamber to the bottom five outlets are nearly identical on both sides. Fluid momentum arriving at the two chamber boundaries remains balanced, resulting in two velocity peaks of comparable magnitude and a symmetric velocity profile. In contrast, the Tesla-valve configuration yields an asymmetric velocity distribution, where the left peak is distinctly lower than the right peak.

This asymmetry arises from two coupled factors: the directional hydraulic resistance inherent to Tesla valves, and the asymmetric layout of the full flow paths extending from the inlet to the bottom row of five outlets. The Tesla valve upstream of the left-side horizontal narrow connecting channels sits left of the chip’s vertical symmetry axis, forming a longer overall flow-path toward its corresponding bottom outlet. Fluid travels through this left Tesla valve in the reverse direction; Tesla valves exhibit high hydraulic resistance under reverse flow, which further elevates the total resistance of this left branch. By comparison, the Tesla valve upstream of the right-side horizontal narrow connecting channels is positioned exactly on this central axis, yielding a shorter flow-path to the bottom outlet. With a fixed total inlet flow rate, the higher total resistance on the left side restricts the flow delivered to the left boundary of the second chamber, reducing the magnitude of the left velocity peak. Meanwhile, the shorter, lower-resistance right-side pathway permits greater fluid momentum to reach the right chamber boundary, forming a higher velocity peak. This asymmetric velocity feature persists across all tested inlet flow rates, confirming that it is an inherent structural effect originating from the combined influence of path-length-dependent hydraulic resistance and the anisotropic flow resistance of Tesla valves. Overall, compared with the conventional curved channel, the Tesla-valve architecture slightly suppresses the bulk flow velocity inside the chamber.

Beyond velocity regulation, the Tesla-valve channel achieves superior performance in generating steady concentration gradients (Fig. S3). At the high flow rate of 30 μL/min, the two concentration profiles nearly coincide near the chamber inlet; as fluid propagates rightward, the Tesla-valve channel declines more steeply through the mid-chamber region but retains a higher residual concentration on the right side. At the moderate flow rate of 10 μL/min, the profiles exhibit a double-crossing behavior: the Tesla-valve channel starts with a marginally higher concentration near the left inlet, falls below the conventional curved channel across the central portion of the chamber, and rises above it again toward the right boundary. Under the low-flow condition (1 μL/min), the Tesla-valve profile begins below the conventional curved channel and gradually overtakes it at the right side of the chamber, with both curves showing continuous concentration decay. Collectively, across all tested flow rates, the concentration profiles of the Tesla-valve channel exhibit smoother and more gradual transitions, delivering a wider and more stable concentration gradient window compared with the conventional curved flow channel.
